# Unraveling APA dynamics during quiescence transitions reveals Pabp modulation of *sod2* expression via APA to regulate cellular quiescence

**DOI:** 10.64898/2026.09.27.754748

**Authors:** Xiaoling Deng, Lei Zhang, Zhirui Du, Xianjia Zhao, Bandhan Sarker, Yaruq Jabeen, Lili Guan, Tianjiao Zhou, Lianming Liang, Shuaishuai Huang, Yonghong Zhou, Chuan Xu

## Abstract

Quiescence is a reversible proliferation arrest enabling cells to adapt to environmental changes, yet the post-transcriptional mechanisms governing its transitions remain poorly understood. Here, we map genome-wide alternative polyadenylation (APA) dynamics across 15 time points of quiescence transitions in fission yeast using 3’-end RNA sequencing. We reveal that APA undergoes extensive remodeling: driving global 3’UTR lengthening during early quiescence entry, 3’ UTR shortening upon quiescence establishment, and progressive reversal to proliferative-state lengths during quiescence exit. Integrating 3’ UTR dynamics with differential expression analysis, we identify candidate genes potentially regulated through APA-mediated 3’ UTR mechanisms. We demonstrate that *sod2*, encoding mitochondrial superoxide dismutase, autoregulates its mRNA and protein levels via APA-dependent 3’ UTR changes. Disrupting proximal APA site of *sod2* in a mitophagy-deficient (*atg43*Δ) background exacerbates quiescence maintenance defects and mitochondrial dysfunction. We further identify Pabp as an upstream regulator of *sod2* APA. This study establishes the temporal APA remodeling during quiescence transitions and reveals a Pabp*-*APA*-sod2* regulatory axis governing post-transcriptional control of cellular quiescence.

## INTRODUCTION

Quiescence (G0) is a highly conserved survival strategy across eukaryotes, from yeast to humans, enabling cells to reversibly arrest proliferation under adverse conditions while retaining the capacity to resume growth upon environmental improvement [1, 2]. This reversible nature distinguishes quiescence from terminal differentiation and cellular senescence [3]. In unicellular organisms, quiescence ensures survival during nutrient exhaustion [4]; in multicellular organisms, it maintains the long-term functional reserve of adult stem cells, including hematopoietic stem cells, hair follicle stem cells, and hepatocytes [5]. Dysregulation of quiescence has been implicated in tissue ageing, regenerative failure, and tumor drug resistance [5–10]. Despite its recognized importance, the regulatory mechanisms governing quiescence transitions remain incompletely understood.

In eukaryotic cells, quiescence transitions are accompanied by global transcriptional repression and chromatin remodeling. These changes encompass redistribution of RNA polymerase II at promoter regions, histone hypoacetylation, redistribution of H3K9me3, and the involvement of chromatin regulators such as HIRA and INO80 [11–15]. For instance, the HIRA complex deposits the histone variant H3.3 during quiescence to maintain heterochromatin integrity and prevent genomic instability [16], whereas the INO80 remodeling complex modulates nucleosome occupancy and dynamics to facilitate RNA polymerase II redistribution and promoter-proximal pausing, thereby fine-tuning transcriptional potential [11]. However, these studies have focused predominantly on transcriptional regulation, and the post-transcriptional control of quiescence transitions, particularly the fine regulation of mRNA maturation, remains relatively unexplored.

Alternative polyadenylation (APA), a paradigmatic post-transcriptional regulatory mechanism, modulates RNA length, particularly 3’ UTR length, through the selection of distinct polyadenylation sites (PAS), thereby influencing mRNA stability, translational efficiency, and subcellular localization to ultimately shape gene expression output [17, 18]. APA is pervasive in multicellular eukaryotes; for example, over 70% of human genes undergo APA [19], and it plays critical regulatory roles in cell proliferation, differentiation, development, immunity, and disease [17]. APA is equally prevalent in unicellular fungi: more than 80% of mRNA genes in *Saccharomyces cerevisiae* (*S. cerevisiae*) and *Schizosaccharomyces pombe* (*S. pombe*) exhibit APA [20], and APA contributes to microbial stress adaptation [21]. Although a limited number of studies have begun to reveal roles for APA in quiescence regulation [22], whether APA undergoes dynamic changes throughout the entire quiescence transition, what the temporal patterns of such changes are, and whether they serve specific biological functions remain largely unknown.

Here, using fission yeast as a model system, we systematically investigate the temporal dynamics of APA during quiescence transitions and the mechanisms by which APA modulates gene expression to influence quiescence states. By manipulating nitrogen availability in the culture medium, we recapitulated the full quiescence cycle. Through mRNA 3’-end transcriptome sequencing and analysis at 15 time points, we constructed a genome-wide APA dynamic landscape during quiescence transitions, revealing the prevalence and genomic features of APA in fission yeast. By integrating APA dynamics with gene expression changes, we identified multiple quiescence-associated genes whose expression is potentially regulated by APA. Among these, we focused on *sod2*, encoding mitochondrial superoxide dismutase, and discovered a novel pathway in which Pabp regulates *sod2* expression through APA-mediated 3’ UTR remodeling of *sod2* to influence quiescence. Our findings highlight the critical role of post-transcriptional regulation in quiescence adaptation and stress tolerance, providing an important case study for understanding post-transcriptional control of cellular quiescence transitions.

## RESULTS

### Pervasive APA in *S. pombe*

We used the heterothallic (*h^-^*^S^) prototrophic fission yeast cells as the experimental system to construct a comprehensive alternative polyadenylation (APA) landscape [4, 23]. Entry into quiescence was induced by nitrogen-source deprivation, whereas exit from quiescence was induced by nitrogen replenishment [23]. Based on previous studies of quiescence [4, 24, 25], we designed 15 sampling points (**Fig. 1A**), including the vegetative growth stage (VE); nine time points during quiescence entry, corresponding to 0.5, 1, 2, 3, 4, 8, 16, 24 and 48 h after nitrogen removal (RN); and five time points during quiescence exit, corresponding to 1, 2, 3.5, 6 and 9 h after nitrogen addition (AN). Two independent biological replicates were collected for each time point.

**Fig. 1.**
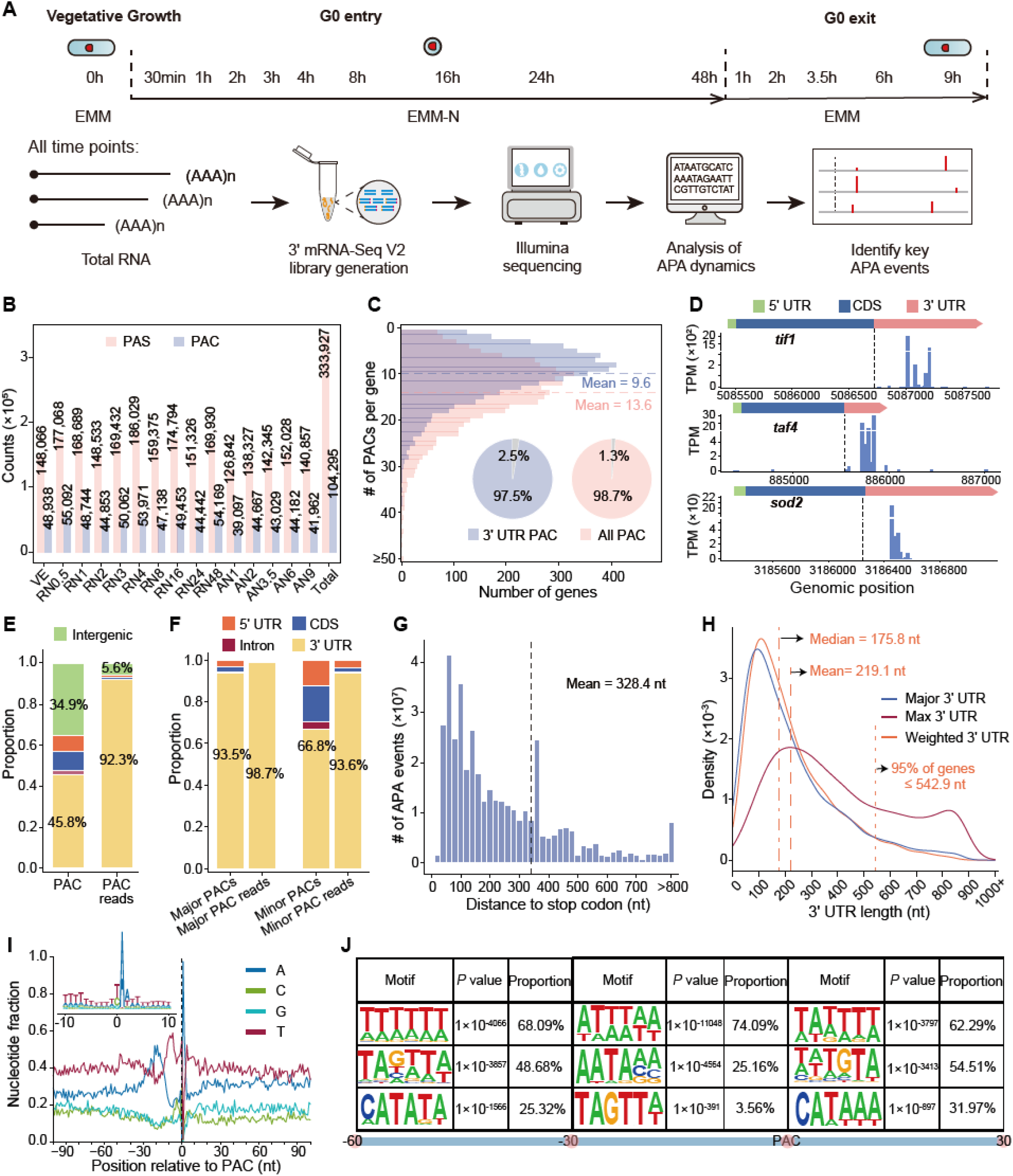
Genomic characteristics of APA in *S. pombe*. (**A**) Schematic of the study design. Fission yeast cells were sampled across 15 time points during quiescence transition, including vegetative growth (VE), nitrogen starvation (EMM-N, G0 entry) at 0.5 h (RN0.5), 1 h (RN1), 2 h (RN2), 3 h(RN3), 4 h (RN4), 8 h (RN8), 16 h (RN16), 24 h (RN24) and 48 h (RN48), and nitrogen replenishment (G0 exit) at 1 h (AN1), 2 h (AN2), 3.5 h (AN3.5), 6 h(AN6) and 9 h (AN9) following 48 h of starvation. (**B**) Total numbers of PASs and PACs detected across the fifteen time points. (**C**) Distribution of gene counts based on the number of expressed PACs and 3’ UTR PACs. Dashed lines represent the mean number of PACs per gene. Insets show the proportion of APA genes. (**D**) Examples of genes producing multiple PACs located in different genomic regions. (**E**) Distribution of PACs and PAC reads across different genomic regions. (**F**) Distribution of major and minor PACs and PAC reads across different genomic regions. (**G**) Distance between PACs and the stop codon, weighted by expression abundance. (**H**) Length distribution of 3’ UTRs determined using three methods. “Major 3’ UTR” refers to the 3’ UTR of the major PAC; “Max 3’ UTR” represents the 3’ UTR calculated from the most distal PAC; and “Weighted 3’ UTR” indicates the average 3’ UTR length calculated from all PACs downstream of the stop codon, weighted by their respective expression abundances. (**I**) Nucleotide distribution in the flanking regions of PAC. (**J**) Top 3 motifs enriched in the −60 to −31, −30 to −1, and 0 to +30 nt regions relative to PACs.

Cells were collected at each time point, and total RNA was extracted for library preparation using Quant-seq 3’-end mRNA-seq (REV) (see **MATERIALS AND METHODS**). In total, 30 sequencing libraries were generated, yielding more than 484 million 3’-end sequencing reads (**Tab. S1**). Following read-quality control and removal of internal-priming events (see **MATERIALS AND METHODS**), we obtained more than 456.8 million mRNA 3’-end reads (**Tab. S1**). To identify high-confidence polyadenylation sites (PASs), we required at least four supporting reads, a more stringent threshold than the two-read threshold used in previous studies [26, 27]. This procedure identified 333,927 PASs, with 126,842-186,029 PASs detected per sample **(Fig. 1B**; **Tab. S1**). Correlation analysis between biological replicates showed that the expression levels of PASs were highly reproducible across all conditions, with correlation coefficients exceeding 0.87 (range, 0.879-0.994), demonstrating the robustness and reliability of the dataset (**Fig. S1A**).

Because APA is intrinsically heterogeneous, we clustered nearby PASs into PAS clusters (PACs). The genomic position of each PAC was defined by the most highly expressed PAS within that cluster, whereas the read count of each PAC was calculated as the sum of reads supporting all PASs in the cluster (see **MATERIALS AND METHODS**). In total, we identified 104,295 PACs, with 39,097-55,092 PACs detected per sample (**Fig. 1B; Tab. S1**). Saturation analyses performed using the combined dataset and separately for each condition showed that the number of detected PACs approached saturation in all cases, indicating that the sequencing depth and sampling scheme were sufficient (**Fig. S1B**).

To evaluate the reliability of our PACs, we compared them with previously reported fission yeast APA datasets from Liu et al. [20]. We found that 33,263 of our PACs (31.9%) were supported by the external datasets (**Fig. S1C**). Although the remaining 71,389 PACs were not supported by the external dataset, 67,954 of them were supported by at least two of our own samples. Consequently, a total of 97% of our PACs were supported by either the external dataset or multiple samples in the present dataset (**Fig. S1C**). While the final 3,078 PACs (3%) were sample-specific, they were still supported by at least 4 reads in corresponding samples. To further validate the reliability of our APA sites, we performed 3’ RACE assays on several example genes (see **MATERIALS AND METHODS**). As illustrated in **Figs. S1D-S1F**, the majority of the sequencing-detected PACs were successfully confirmed by PCR amplification and gel electrophoresis. Collectively, these results demonstrate the high confidence of our PACs.

We next examined the number of PACs per gene. On average, each *S. pombe* gene expressed 13.6 PACs, and 98.7% of genes expressed more than one PAC (**Fig. 1C**). This proportion is higher than those reported for humans (67%) [28, 29], mice (56%) [26], zebrafish (43%) [30] and *Arabidopsis thaliana* (>90%) [31]. Nevertheless, such comparisons should be interpreted with caution because APA sequencing strategies, data-processing procedures and the intrinsic properties of APA differ substantially among species, limiting direct cross-species comparisons. Even considering PACs only located within 3’ UTRs, each *S. pombe* gene expressed an average of 9.6 PACs, and 97.5% of genes expressed more than one 3’ UTR PAC (**Fig. 1C**). For example, 18 PACs were detected within the 3’ UTR of *tif1* across all time points (**Fig. 1D**). Four PACs detected for *tif1* under VE conditions were independently validated by 3’ RACE (**Fig. S1D**). In addition, PACs detected under VE conditions for *taf4* and *sod2* were also confirmed by 3’ RACE (**Figs. S1E and S1F**).

Notably, the number of identified PACs is highly sensitive to analytical parameters. Specifically, greater sequencing depth and a larger number of samples facilitate the detection of additional PACs (**Fig. S1B**). Similarly, applying more permissive thresholds for cluster width or the minimum number of supporting reads also increases the number of defined PACs (**Fig. S1G**). In this study of *S. pombe*, we adopted the cluster width threshold from a previous APA analysis by Liu et al. [20] to ensure comparability with prior research and with *S. cerevisiae*. However, given our higher sequencing coverage, we applied a more stringent threshold for supporting reads of a PAS to ensure the accurate identification of genuine PACs.

To provide a comprehensive view of the fission yeast APA landscape, we compiled our genome-wide APA data together with selected datasets from Liu et al. [20] to construct spoAPADb, a publicly accessible database dedicated to querying and disseminating APA sites in *S. pombe* (available at: https://gpd.sjtu.edu.cn/spoapadb/).

### Genomic characteristics of APA in *S. pombe*

Leveraging our high-coverage dataset than before [20], we next briefly investigated the genomic features of APA in *S. pombe*. Because most *S. pombe* genes lack annotated UTRs, we extended gene boundaries using public RNA-seq data following a previously established method [20] (see **MATERIALS AND METHODS**). Genomic mapping revealed that although 34.9% of PACs reside in intergenic regions, they account for only 5.6% of PAC reads (**Fig. 1E**). Conversely, 3’ UTR PACs constitute 45.8% of all PACs but capture 92.3% of PAC reads (**Fig. 1E**). For instance, PACs for *tif1* and *sod2* were exclusively observed in their 3’ UTRs (**Fig. 1D**). The remaining PACs were distributed across CDS (9.5%, 0.8% reads), 5’ UTR (7.7%, 1.0% reads), and introns (2.1%, 0.16% reads) (**Fig. 1F**). For example, *taf4* harbored PACs in both its CDS and 5’ UTR (**Fig. 1D**). This predominant 3’ UTR localization is consistent with the canonical role of APA in transcription termination.

We further classified PACs in a gene into major (i.e., the most abundant one) and minor PACs [32]. Over 93.5% of major PACs reside in annotated 3’ UTRs, accounting for 98.7% of PAC reads (**Fig. 1F**), underscoring their essential role in canonical gene expression. Although minor PACs are also predominantly 3’ UTR-localized (66.8% of PACs, 93.6% of reads), they show substantial representation in CDS, introns, and 5’ UTRs (**Fig. 1F**). This broader distribution suggests that minor PACs may contribute to transcript diversity and fine-tuned transcriptional regulation [33, 34].

We then defined 3’ UTR length as the distance from the annotated stop codon to downstream PACs. Transcript-level 3’ UTR lengths in *S. pombe* ranged predominantly from 40 to 500 nt, with a mean of 328.4 nt (**Fig. 1G**). At the gene level, we calculated three metrics: weighted average, major PAC-associated, and most distal PAC-associated 3’ UTR lengths (see **MATERIALS AND METHODS**; **Fig. 1H**). Based on the weighted metric, the mean and median 3’ UTR lengths were 219.1 nt and 175.8 nt, respectively, with 95% of genes having 3’ UTRs shorter than 542.9 nt (**Fig. 1H**).

Sequence context analysis revealed distinct nucleotide patterns surrounding APA sites (**Fig. 1I**). The region from-100 to −30 nt upstream is TA-rich, followed by an A-rich zone (−30 to −15 nt) and a T-rich zone (−15 to −1 nt). The PAC site itself (+0 to +3) is highly A-enriched, whereas the downstream region (+4 to +100 nt) is predominantly T-rich. Top enriched 6-mer motifs further highlighted these patterns (**Fig. 1J**): TTTTTT, TAGTTA, and CATATA dominate the −60 to −31 nt region; ATTTAA and AATAAA are enriched from −30 to −1 nt; and TATTTT, TATGTA, and CATAAA are prevalent downstream (0-30 nt). Notably, the canonical mammalian polyadenylation signal AATAAA [35] is significantly enriched in *S. pombe*, suggesting a conserved APA mechanism between *S. pombe* and mammals [20, 36].

### Global APA remodeling coordinates with quiescence transitions

To compare APA patterns across different time points during quiescence transition, we first addressed potential biases caused by varying sequencing depths. We down-sampled the uniquely mapped PAC reads for each time point to an equal depth of 22.3 million reads and re-evaluated the genomic distribution and abundance of PACs (see **MATERIALS AND METHODS**). Principal component analysis (PCA) of PAC expression levels clearly separated the quiescence entry and exit groups. Notably, the AN9 sample clustered most closely with the VE stage (**Fig. 2A**), indicating that PAC expression profiles accurately reflect cellular states and further validating the robustness of our dataset.

**Fig. 2.**
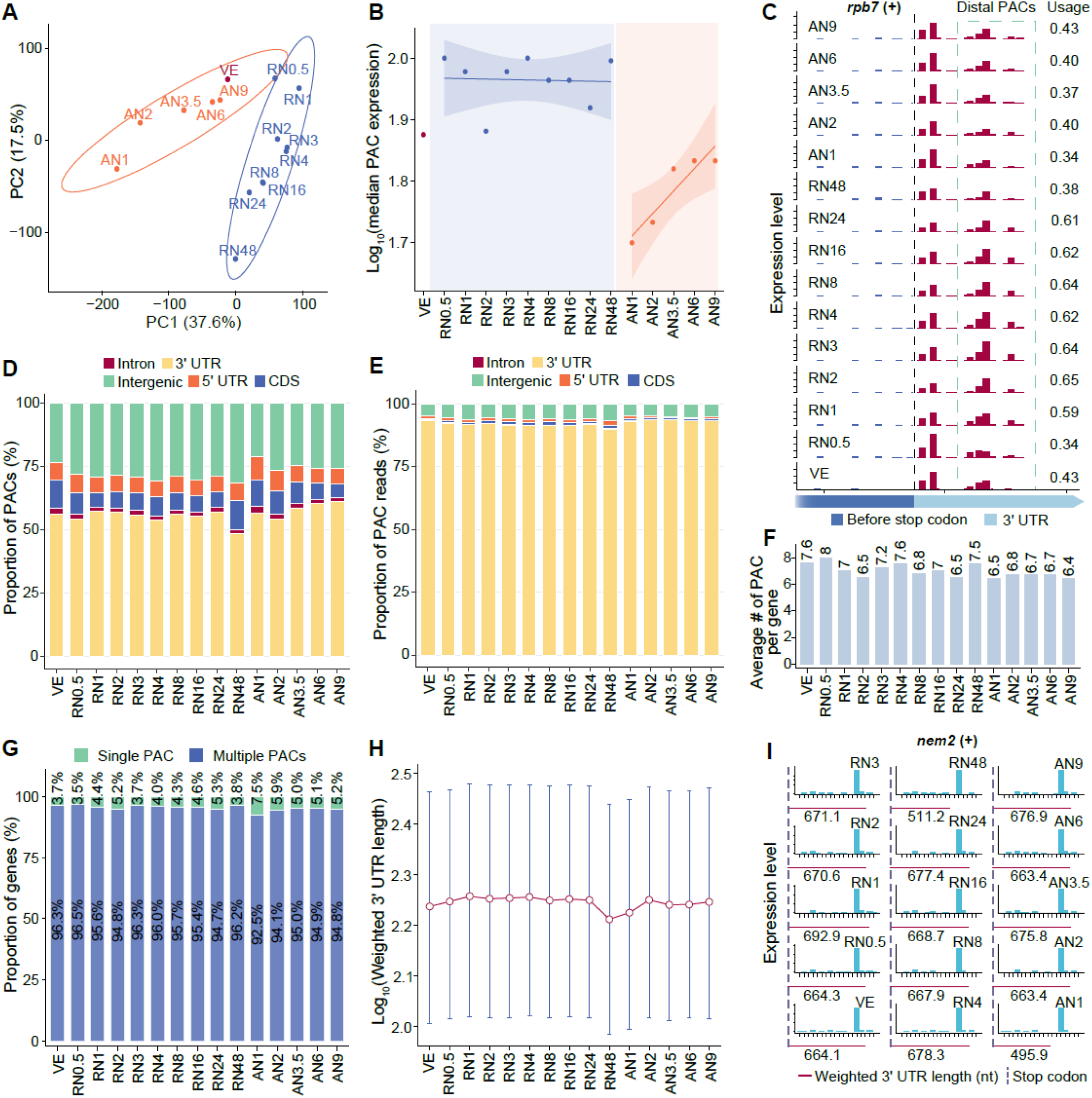
Dynamical global remodeling of APA during quiescence transitions. **(A)** PCA analysis of PAC expression across different time points. **(B)** Dynamic changes in PAC expression over time points. **(C)** Examples of gene with different APA profiles in 15 time points. (**D**) Distribution of PACs across different genomic regions in various time points. (**E**) Distribution of PAC reads across different genomic regions in various time points. (**F**) Average number of PACs per gene across time points. (**G**) Proportion of genes with single or multiple PACs in 15 time points. (**H**) Distribution of weighted 3’ UTR length across different time points. (**I**) Representative example of gene with weighted 3’ UTR length change across 15 time points.

Using the median PAC expression to represent overall activity, we observed distinct differences among the VE, quiescence entry, and exit stages. PAC expression was consistently higher during quiescence entry than in the VE stage, whereas it was lower across all time points during quiescence exit (**Fig. 2B**). During the exit stages, PAC expression was initially at its lowest but gradually increased over time, approaching VE levels (**Fig. 2B**). For example, the distal PAC of the *rpb7* gene exhibited high expression throughout quiescence entry, but its expression markedly decreased to VE levels during the exit stages (**Fig. 2C**).

We next examined the genomic distribution of PACs and found both the number (**Fig. S2A**) and proportion (**Fig. 2D**) of intergenic PACs increased during the transition from VE to quiescence entry. Conversely, during the transition from quiescence to the exit stages, intergenic PACs significantly decreased, gradually recovering to near-VE levels (**Figs. S2A and 2D**). This trend was mirrored when analyzing the distribution and proportion of PAC reads (**Figs. 2E and S2B**).

Within genic regions, the proportion of 3’ UTR PACs reached its nadir at 48 hours of nitrogen starvation (RN48, fully quiescent), although the absolute number of PACs was lowest at 1 hour of nitrogen replenishment (**Figs. S2A and 2D**). In terms of PAS reads, both the count and proportion of 3’ UTR PACs decreased during quiescence entry, reaching a minimum at RN48, and subsequently increased during the exit stages (**Figs. 2D and S2B**). These distribution patterns indicate systematic APA remodeling during quiescence transitions.

We also calculated the average number of PACs per gene at each time point (**Figs. 2F and S2C**). While no clear trend was observed when considering all genomic PACs (**Fig. 2F**), the number of 3’ UTR PACs per gene dropped to a minimum of 4.8 at 1 hour of nitrogen replenishment and then progressively recovered toward VE levels (**Fig. S2C**). A similar pattern was observed for genes with multiple PACs, where both the number and proportion of such genes were lowest at 1 hour of nitrogen replenishment and gradually increased during quiescence exit (**Figs. 2G, S2D, and S2E**).

To assess global 3’ UTR length dynamics, we calculated the weighted 3’ UTR length for each gene based on the distance between the stop codon and downstream PACs (**Fig. 2H**). Compared to the VE stage, global 3’ UTR elongation occurred during quiescence entry, followed by significant shortening at RN48. During quiescence exit, 3’ UTR lengths showed a tendency to recover toward VE levels. For instance, the *nem2* gene exhibited increased distal APA usage and 3’ UTR elongation during quiescence entry stages, significant shortening at RN48, and a gradual lengthening back toward VE levels during quiescence exit stages (**Fig. 2I**).

In summary, while global APA profiles remained relatively similar during the early stages of quiescence entry, distinct APA patterns emerged upon the full entry into quiescence (e.g., at 48 hours). These changes were reversed during the quiescence exit stages, with APA profiles recovering to near-VE states. Thus, APA remodeling is tightly coordinated with the physiological transitions associated with quiescence.

### Substantial alterations in 3’UTRs induced by tandem APAs during quiescence transitions

We next investigated 3’ UTR length dynamics driven by tandem APAs in 3’ UTR region. To quantify these changes during quiescence transition, we calculated the Percentage of Distal polyA site Usage Index (PDUI) (**Figs. 3A and S3A**). Genes with significantly increased PDUI values were classified as undergoing 3’ UTR lengthening, those with decreased PDUI as shortening, and those without significant changes as unchanged (see **MATERIALS AND METHODS**).

**Fig. 3.**
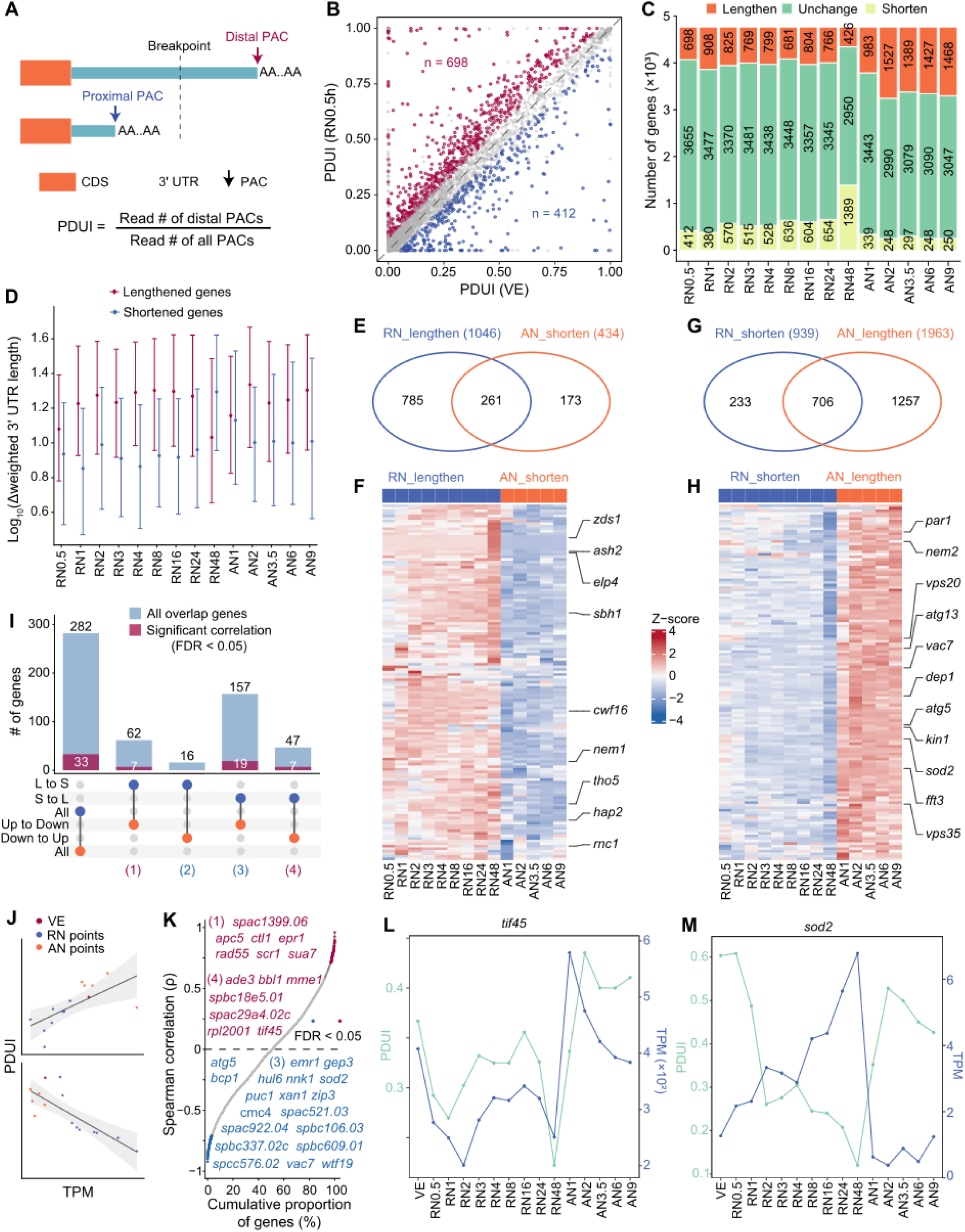
3’UTR alterations induced by tandem APAs during quiescence transitions. (**A**) Schematic illustrating the definition of proximal and distal PACs. (**B**) Representative example (RN0.5) showing the number of genes with altered 3’ UTR length. (**C**) Quantification of genes with altered 3’ UTR length in each time point. (**D**) Range of Δ3’ UTR length across time points, stratified by genes undergoing 3’ UTR lengthening and shortening. (**E**) Overlap of genes with 3’ UTR lengthening in quiescence entry stages and 3’ UTR shortening in quiescence exit stages. (**F**) Heatmap showing ΔPDUI values for genes exhibiting 3’ UTR shortening and lengthening during across 14 time points during quiescence transitions. (**G**) Overlap of genes with 3’ UTR shorten in quiescence entry stages and 3’ UTR lengthen in quiescence exit stages. (**H**) Heatmap showing ΔPDUI values for genes exhibiting 3’ UTR lengthening and shortening across 14 time points during quiescence transitions. In (**F**) and (**H**), ΔPDUI values were calculated relative to VE for quiescence entry and rn48 for quiescence exit. Red indicates increased distal poly(A)-sites usage (3’ UTR lengthening), whereas blue indicates decreased distal poly(A)-sites usage (3’ UTR shortening). Selected quiescence-related genes are labeled on the right. **(I)** Overlap of genes with both the PDUI and expression changed, classified into 4 types. **(J)** Schematic illustrating positive and negative related between PDUI and gene expression levels (TPM). **(K)** Spearman correction between PDUI and TPM of all genes. Significant genes are marked. **(L)** and **(M)** show profiles of PDUI and TPM of two gene examples.

During quiescence entry stages, we compared each time point against the VE stage. For example, at the RN0.5 time point, 14.6% (698 genes) exhibited significantly higher PDUI values than in the VE stage, whereas only 8.6% (412 genes) showed significantly lower PDUI values (**Fig. 3B**). This significant predominance of 3’ UTR lengthening at the RN0.5 stage aligns with the genome-wide 3’ UTR shifts observed in **Fig. 2H**. We extended this analysis across all nine entry time points and found that only a minority of genes exhibited altered 3’ UTR lengths (**Fig. 3C**). Nevertheless, during the first eight time points, the number of genes with lengthened 3’ UTRs consistently exceeded those with shortened 3’ UTRs. Conversely, at the RN48, the number of genes with shortened 3’ UTRs surpassed those with lengthened ones (**Fig. 3C**), further corroborating the genome-wide trends shown in **Fig. 2H**.

Beyond the RN48 time point, the magnitude of 3’ UTR lengthening generally exceeded that of shortening (**Fig. 3D**), consistent with the observation that global 3’ UTR lengths are shortest at RN48 (**Fig. 2H**). Since RN48 represents the full entry into quiescence, this suggests that the distinct physiological state of quiescence is reflected in specific APA signatures.

For the quiescence exit stages, we used the RN48 time point as a baseline to investigate APA-driven 3’ UTR changes across five exit time points. We observed a gradual lengthening of 3’ UTRs during exit stages; specifically, compared to RN48, the majority of genes with altered 3’ UTRs exhibited lengthening at all exit time points, with over 1,000 genes affected at each point (**Fig. 3C**). Furthermore, the magnitude of lengthening consistently exceeded that of shortening among these genes (**Fig. 3D**). These findings indicate that the 3’ UTR shortening characteristic of full quiescence entry is reversed during the exit stages.

We then defined genes with 3’ UTR changes associated with quiescence entry as those showing significant differences relative to the VE stage in at least one time point, without exhibiting significant opposing changes at any other entry time point. Using this criterion, we identified 1,985entry-associated genes with altered 3’ UTR lengths, comprising 1,046 lengthened (**Fig. 3E**) and 939 shortened genes (**Fig. 3G**). Similarly, exit-associated genes were defined as those showing significant differences relative to RN48 in at least one time point, without significant opposing changes at any other exit time point. This yielded 2,397 exit-associated genes, including 1,963 lengthened (**Fig. 3G**) and 434 shortened genes (**Fig. 3E**).

Given that quiescence entry and exit are reciprocal processes, genes exhibiting opposing 3’ UTR changes between these two stages are likely most relevant to the quiescence transition. Venn diagram analysis revealed 261 genes that were lengthened during quiescence entry but shortened during quiescence exit (**Fig. 3E**). Examples include *tho5*, *sbh1*, and *cwf16* (**Fig. 3F**), which have been previously linked to quiescence [37]. Conversely, 706 genes were shortened during quiescence entry but lengthened during quiescence exit (**Fig. 3G**), including *par1*, *vac7* and *atg5* (**Fig. 3H**), also known to be associated with quiescence [37].

Gene Ontology (GO) term enrichment analysis showed that genes lengthened during quiescence entry and shortened during quiescence exit are primarily enriched in molecular functions such as RNA 2’-O-methyltransferase activity, protein folding chaperone activity, and membrane bending activity, as well as biological processes including mRNA splicing via spliceosome, RNA splicing, and endoplasmic reticulum organization (**Fig. S3B**). In contrast, genes shortened during quiescence entry and lengthened during quiescence exit are enriched in molecular functions like cysteine-type endopeptidase activity, translation initiation factor binding, and glycosyltransferase activity, alongside cellular components and biological processes involving the proteasome complex, spindle microtubules, and deactivation of the mitotic spindle assembly checkpoint (**Fig. S3B**). Notably, both gene sets are enriched in biological processes such as regulation of translation, regulation of biosynthetic processes, regulation of primary metabolic processes, and post-transcriptional regulation of gene expression. This suggests that oppositely directed 3’ UTR dynamics may coordinately contribute to the remodeling of protein synthesis and cellular metabolic states during quiescence transitions.

### Coupling between gene expression and 3’UTR alterations during quiescence transitions

Previous studies have reported that APA can modulate gene expression levels by altering 3’ UTR lengths [38, 39]. We therefore investigated whether the observed 3’ UTR dynamics might regulate transcriptional abundance during quiescence transitions. To this end, we first performed differentially expressed gene (DEG) analysis across the quiescence transitions (see **MATERIALS AND METHODS**).

For the nine quiescence entry time points, we compared each against the VE stage. At the RN0.5 time point, for example, we identified 574 upregulated and 196 downregulated genes (**Fig. S4A**). Across all nine entry time points, the number of upregulated genes ranged from 574 to 1,300, while downregulated genes ranged from 186 to 528 (**Fig. S4B**). For the five quiescence exit time points, we used the RN48 time point as the baseline for comparison (**Fig. S4B**). Across these exit time points, upregulated genes ranged from 451 to 619, and downregulated genes ranged from 1,143 to 1,661.

We defined quiescence-associated DEGs as those showing significant differential expression in at least one time point without exhibiting significant opposing changes at any other time point within the same quiescence phase. Using this criterion, we identified 2,052 upregulated (**Fig. S4C**) and 967 downregulated (**Fig. S4D**) genes associated with quiescence entry. For quiescence exit, we identified 956 upregulated (**Fig. S4C**) and 2,290 downregulated (**Fig. S4D**) genes.

We next focused on genes exhibiting opposing expression patterns between quiescence entry and exit. Venn diagram analysis revealed an overlap of 1,497 genes that were upregulated during entry and downregulated during exit (**Fig. S4C**). Examples include *moh1*, *cqd1*, and *iec1* (**Fig. S4E**), which have been previously linked to quiescence [37]. Conversely, 520 genes were downregulated during entry but upregulated during exit (**Fig. S4D**), including *SPBC365.16*, *lys12*, and *maa1* (**Fig. S4F**), also known to be associated with quiescence [37].

To identify genes whose expression levels might be potentially regulated by 3’ UTR remodeling during quiescence transitions, we integrated our 3’ UTR dynamics data with the DEG analysis. We found an overlap of 282 genes that exhibited both 3’ UTR length changes and significant expression changes during the transition, representing candidate targets for 3’ UTR-mediated regulation (**Fig. 3I**). Because 3’ UTR length variations can either upregulate or downregulate gene expression depending on the specific gene [39–42], we further categorized these 282 genes into four classes based on their 3’ UTR and expression changes during quiescence entry: (1) 3’ UTR lengthening with upregulated expression (corresponding to 3’ UTR shortening with downregulated expression during quiescence exit); (2) 3’ UTR lengthening with downregulated expression (corresponding to 3’ UTR shortening with upregulated expression during quiescence exit); (3) 3’ UTR shortening with upregulated expression (corresponding to 3’ UTR lengthening with downregulated expression during quiescence exit); and (4) 3’ UTR shortening with downregulated expression (corresponding to 3’ UTR lengthening with upregulated expression during quiescence exit). We identified 62, 16, 157, and 47 genes in classes 1, 2, 3, and 4, respectively (**Fig. 3I**).

To further validate the relationship between 3’ UTR length and gene expression, we calculated the Spearman rank correlation between PDUI values and expression levels for these candidate genes. A positive correlation was defined as positive regulation, whereas a negative correlation indicated negative regulation (**Fig. 3J**). We identified fourteen genes with significant positive correlations (**Fig. 3K**), belonging to classes 1 (7 genes) and 4 (7 genes) (**Fig. 3I**), suggesting that distal APA site usage positively regulates their expression. Additionally, 19 genes showed significant negative correlations (**Fig. 3K**), all belonging to class 3 (**Fig. 3I**), indicating that distal APA site usage negatively regulates their expression. For example, *tif45*, which encodes the translation initiation factor eIF4E and is involved in the initiation and regulation of protein synthesis [43], exhibited a significant positive correlation between its PDUI and expression level (**Fig. 3L**). In contrast, the *sod2* gene showed a significant negative correlation between its PDUI and expression level (**Fig. 3M**). These findings provide a set of target genes for our subsequent functional and mechanistic investigations.

### Reciprocal dynamics of distal APA usage and gene expression of *sod2* during quiescence transitions

To validate our above integrated analysis, we selected *sod2*, which encodes mitochondrial superoxide dismutase [44] and has been previously implicated in quiescence [37], for experimental validation. We designed two pairs of specific RT-qPCR primers to quantify distal PACs (dPACs) usage (**Fig. 4A**). The S-F/S-R primer pair amplifies transcripts from both proximal PACs (pPACs) and dPACs (total *sod2* mRNA), whereas the L-F/L-R pair specifically amplifies dPAC-containing transcripts. The ratio of L-F/L-R to S-F/S-R product yields the PDUI of *sod2* gene. Quantitative analysis across time points (**Fig. 4B**) showed that dPAC usage is high in the VE stage, corresponding to the longer 3’ UTR isoform. During quiescence entry, PDUI gradually decreased, indicating a shift toward the shorter 3’ UTR isoform, with the lowest PDUI observed at RN48. This trend was reversed during quiescence exit, as PDUI gradually increased to levels comparable to the VE stage. These RT-PCR-derived PDUI dynamics are highly consistent with our APA-seq results (**Fig. 3M**), confirming the reliability of our sequencing data.

**Fig. 4.**
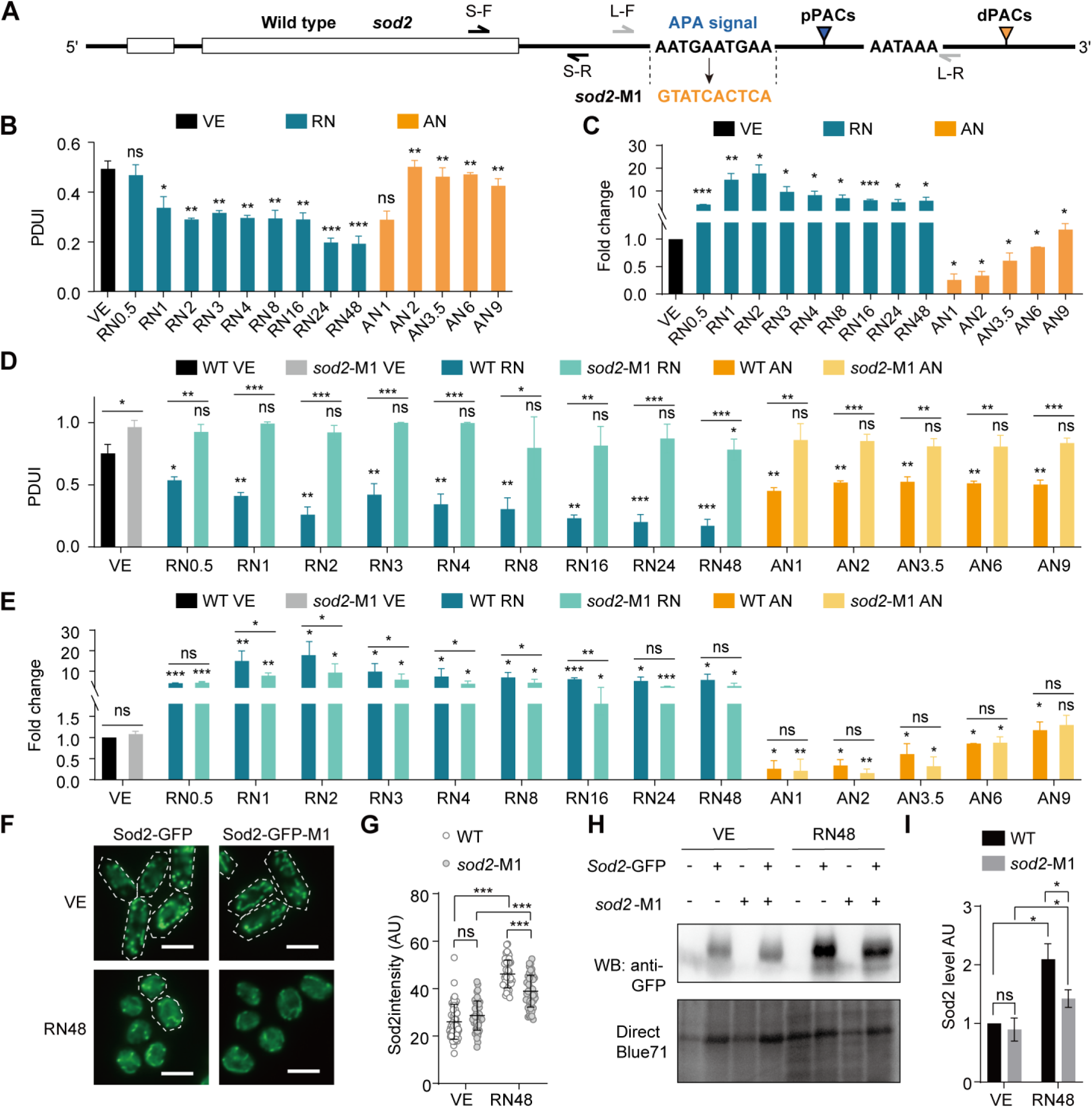
*sod2* APA regulates its expression level. (**A**) Specific PCR primers were designed to quantify the expression of the long and short 3’ UTR isoforms of *sod2*. The nucleic acid sequences of the APA signal in WT cells and *sod2-*M1 were present. (**B**) PDUI and (**C**) expression levels of *sod2* during quiescence transitions. Dark blue bars represent samples during quiescence entry, with statistical significance calculated relative to vegetative growth (VE). Dark yellow bars represent samples during quiescence exit, with statistical significance calculated relative to the 48 h after nitrogen-starvation (RN48). (**D**) PDUI and (**E**) expression levels of *sod2* in WT and *sod2*-M1 cells during quiescence transitions. Black, blue, and yellow bars indicate VE, quiescence entry, and quiescence exit stages, respectively, with dark and light shades denoting WT and *sod2*-M1 strains. For each genotype (WT or *sod2*-M1), statistical significance during entry was determined relative to corresponding VE, and during exit relative to its own 48-h after nitrogen-starvation. Additionally, statistical significance between WT and *sod2*-M1 was determined at each time point. **(F)** Localization of Sod2-GFP in wild-type and Sod2-GFP-M1 cells under VE and RN48 conditions. **(G)** A vertical scatter plot of Sod2-GFP signal intensity in wild-type and Sod2-M1 cells under VE and RN48 conditions. 50 cells were assessed. (**H**) Western blotting of Sod2-GFP in wild-type and Sod2-GFP-M1 cells under VE and RN48 conditions. Protein loading was confirmed by Direct Blue 71 staining. **(I)** The Sod2-GFP band intensity was characterized via three independent western blotting results, and the mean ± S.D. was presented. AU: arbitrary unit. \**P* < 0.05, ** *P* < 0.01 and *** *P* < 0.001 and ns: not significant.

Concurrently, RT-qPCR analysis of gene expression revealed that *sod2* mRNA levels were significantly upregulated at all nine quiescence entry time points relative to the VE stage. During quiescence exit, *sod2* expression was significantly downregulated relative to the RN48 baseline (**Fig. 4C**), gradually increasing over time to reach VE levels by AN9. These expression dynamics also align closely with our RNA-seq data (**Fig. 3M**).

Finally, we examined Sod2 protein levels at the RN48 time point relative to the VE stage. Qualitative GFP imaging demonstrated significantly higher Sod2-GFP signal intensity at RN48 compared to the VE stage (**Figs. S5A and S5B**). Western blotting further confirmed elevated Sod2 protein levels during quiescence (**Figs. S5C and S5D**), consistent with the observed mRNA upregulation (**Fig. 4C**).

### *sod2* APA regulates its expression during quiescence entry

Although the aforementioned results demonstrated a negative correlation between APA-driven 3’ UTR length alterations and *sod2* expression levels during quiescence transitions, they do not definitively establish a causal regulatory role of APA in *sod2* expression, because such coupling could potentially be attributed to other confounding regulatory factors. To elucidate this regulatory mechanism, we subsequently analyzed the sequences flanking the dPACs and pPACs. We identified a tandem AATGAA motif located 20–10 nt upstream of the pPACs, and an AATAAA motif located 42–36 nt upstream of the dPACs (**Fig. 4A**). If APA regulates 3’ UTR length, mutating the upstream AATGAA motif should increase dPAC usage, thereby increasing the PDUI of *sod2*. To test this, we replaced the pPAC-upstream motif with a random sequence to construct the *sod2*-M1 mutant, aiming to impair pPAC usage and enhance dPAC usage (**Fig. 4A**). Indeed, we found that the PDUI of the *sod2*-M1 mutant was significantly higher than that of the wild type (WT) in the VE stage, indicating increased dPAC usage (**Fig. 4D**). Moreover, at nearly all time points, the *sod2*-M1 mutant exhibited a significantly higher PDUI than the WT (**Fig. 4D**), confirming that the replaced motif sequence regulates 3’ UTR lengthening.

Since quiescence entry reduces the PDUI of WT *sod2*, we hypothesized that this decreasing trend would be abolished if the pPAC motif were disrupted. Consistently, we observed no significant difference in the PDUI of *sod2*-M1 between the VE stage and any time point during quiescence entry (**Fig. 4D**). Similarly, during quiescence exit, the PDUI of *sod2*-M1 remained comparable to that at the RN48 time point across all time points (**Fig. 4D**). These results indicate that the impairment of the pPAC motif abolishes the dynamic regulation of *sod2* 3’ UTR length during quiescence transitions, further confirming that APA regulates *sod2* 3’ UTR length.

Because *sod2* expression increases with shorter 3’ UTRs and decreases with longer 3’ UTRs (**Figs. 3M, 4B, and 4C**), and the *sod2*-M1 mutant predominantly uses the longer 3’ UTR isoform throughout quiescence transitions (**Fig. 4D**), we expected lower *sod2* expression in the mutant compared to the WT. Indeed, RT-qPCR results confirmed reduced *sod2* expression during quiescence entry in the mutant (**Fig. 4E**). Interestingly, however, we did not observe a general reduction in *sod2* expression in the *sod2*-M1 mutant during the VE or quiescence exit stages (**Fig. 4E**). This suggests that the regulation of *sod2* expression by APA is context-specific, particularly during quiescence entry, and implies that *sod2* expression is also governed by additional regulatory mechanisms beyond APA.

To investigate the impact of APA-mediated 3’ UTR shortening on protein expression during quiescence entry, we compared Sod2 protein levels between WT and *sod2*-M1 mutant cells at the VE stage and 48 h post-nitrogen starvation (RN48). The RN48h time point was specifically selected because quiescence is fully established by this stage, and *sod2* transcript levels exhibit a downward trend relative to WT (**Fig. 4E**). Qualitative GFP imaging revealed no significant difference in Sod2-GFP signal intensity between the mutant and WT at the VE stage (**Figs. 4F and 4G**). However, at RN48, the Sod2-GFP signal intensity in the mutant was significantly reduced compared to that in the WT (**Figs. 4F and 4G**). These observations were further validated by western blotting (**Figs. 4H and 4I**). Notably, while the mRNA level of *sod2*-M1 was lower than that of WT at RN48h, this difference was not statistically significant (**Fig. 4E**). In contrast, the significant reduction in Sod2 protein levels suggests that the APA-mediated 3’ UTR shortening during quiescence entry may impair translational efficiency, thereby decreasing protein abundance.

### *sod2* APA regulates cellular quiescence under autophagic deficiency

*sod2* has previously been identified as a potential essential gene for quiescence in *S. pombe* [37]; however, its specific function remains unverified. To address this, we first constructed a complete *sod2* deletion strain (*sod2*Δ). Flow cytometry analysis revealed that *sod2*Δ cells can enter quiescence normally (**Fig. S6A**). However, their viability significantly declined during prolonged quiescence maintenance (**Fig. S6B**). Furthermore, flow cytometry indicated that *sod2*Δ cells fail to properly exit quiescence following 48 hours of nitrogen starvation. Specifically, 6 hours after nitrogen replenishment, nearly 50% of WT cells remained at the 1C DNA content stage, whereas all *sod2*Δ cells were arrested at 1C. By 12 hours post-replenishment, the proportion of 1C WT cells dropped to less than 1%, while approximately 20% of *sod2*Δ cells remained trapped at the 1C stage (**Fig. S6A**). Consistently, colony formation assays for detecting the mitotic competence (MC) demonstrated a markedly impaired ability to exit quiescence (**Fig. S6C**), confirming that *sod2* is functionally required for quiescence transitions.

Given that *sod2* expression is partially regulated by APA during quiescence transitions (**Fig. 4**) and *sod2* is a quiescence regulator (**Fig. S6**), we hypothesized that this APA-mediated regulation may play a role in quiescence transitions. To this end, our further experiments found that, after 7 days of starvation and subsequent nitrogen replenishment, approximately 40% of WT cells remained arrested at the 1C stage at 6 hours, whereas only 30% of *sod2*-M1 cells were arrested at 1C, indicating a slightly faster exit from quiescence (**Fig. S6D**). By 12 hours post-replenishment, both strains had fully exited quiescence with no significant difference (**Fig. S6D**). Additionally, trypan blue exclusion and colony formation assays showed that disrupting the pPAC of *sod2* did not significantly affect quiescence entry or maintenance (data not shown).

Considering the crucial role of *sod2* in scavenging mitochondrial reactive oxygen species (ROS) and maintaining redox homeostasis [45, 46], we speculated that the regulatory importance of *sod2* APA might become essential under conditions of impaired mitochondrial quality control. Autophagy deficiency has been shown to cause accumulation of dysfunctional mitochondria and elevated ROS in multiple systems [47, 48]. Thus, we utilized a deletion mutant of the mitophagy receptor gene *atg43* as a model for autophagy deficiency [49]. Atg43 localizes to the mitochondrial outer membrane and is upregulated under nitrogen starvation [49]. It anchors isolation membranes to the mitochondrial surface via its Atg8-family-interacting motif (AIM) to initiate mitophagy. We combined *atg43*Δ and *sod2*-M1 and assessed the phenotypes of the double deletion strain. Spot assays assessing proliferative growth revealed that the *sod2*-M1 *atg43*Δ double mutant exhibited more severe growth defects at various temperatures compared to the *atg43*Δ single mutant (**Fig. S6E**).

Next, we monitored quiescence entry via flow cytometry. WT cells successfully entered quiescence, transitioning from a 2C to a 1C DNA content upon nitrogen starvation. In contrast, *atg43*Δ single mutants arrested at the 2C stage (from an initial 4C state), indicating a defect in quiescence entry. The *sod2*-M1 *atg43*Δ double mutant similarly arrested at 2C, showing no exacerbated entry block compared to *atg43*Δ (**Fig. 5A**). After two weeks of nitrogen starvation followed by 12 hours of nitrogen replenishment, neither the *atg43*Δ single mutant nor the double mutant transitioned from 2C to 4C (**Fig. 5A**). Notably, throughout both starvation and recovery, the double mutant consistently exhibited a higher proportion of cells with low forward scatter (FSC) signals compared to *atg43*Δ, suggesting a higher proportion of dead cells (**Fig. 5A**).

**Fig. 5.**
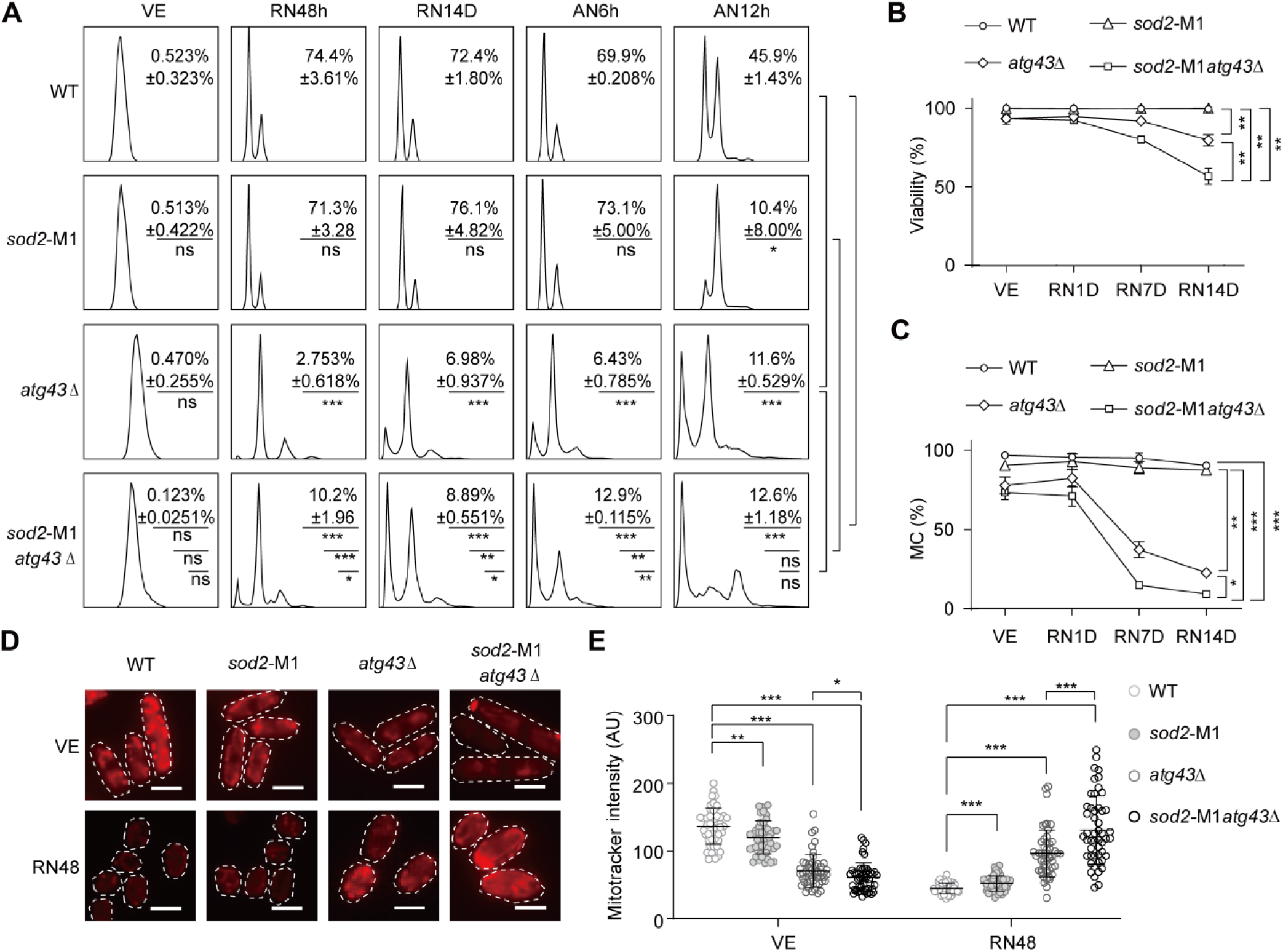
APA of *sod2* is required for cellular quiescence in the absence of *atg43*. (**A**) G0 entry and exit were examined by flow cytometry in WT, *sod2-*M1, *atg43*Δ, and *sod2*-M1 *atg43*Δ cells subjected to nitrogen starvation and subsequent re-addition. Samples were collected during vegetative growth (VE), at 48 h (RN48h) and 14 days (RN14D) after nitrogen depletion and at 6 h (AN6h) and 12 h (AN12h) after nitrogen re-addition. (**B**) Cell viability in WT, *sod2-*M1, *atg43*Δ and *sod2-*M1*atg43*Δ cells following nitrogen starvation. Cell viability was assessed by Trypan blue exclusion analysis, examining over 200 cells per experiment, and data are presented as mean ± S.D. from three independent experiments. (**C**) Colony formation assays evaluated mitotic competence (MC) of WT, *sod2-*M1, *atg43*Δ and *sod2-*M1*atg43*Δ over various time points following nitrogen deprivation. (**D**) localization of MitoTracker red in WT, *sod2-*M1, *atg43*Δ and *sod2-*M1*atg43*Δ cells under vegetative growth and following nitrogen starvation 48h. (**E**) A vertical scatter plot of MitoTracker red signal intensity in WT, *sod2-*M1, *atg43*Δ and *sod2-*M1*atg43*Δ cells under vegetative growth and following nitrogen starvation 48h. 50 cells were assessed. AU: arbitrary unit. * *P* <0.05, ** *P* < 0.01. *** *P* < 0.001.

We further assessed quiescence maintenance. Trypan blue exclusion assays demonstrated that, compared to *atg43*Δ, the *sod2*-M1 *atg43*Δ double mutant suffered a more significant decline in viability during prolonged quiescence (**Fig. 5B**), consistent with the flow cytometry data (**Fig. 5A**). Colony formation assays for detecting MC further confirmed that the double mutant had a reduced capacity to resume proliferation upon exiting quiescence compared to *atg43*Δ (**Fig. 5C**). These results indicate that APA regulation of *sod2* becomes critical for quiescence maintenance and exit specifically under autophagy-deficient conditions.

Given that Sod2 protects mitochondria from oxidative injury [45] and Atg43 mediates the clearance of dysfunctional mitochondria through mitophagy [49], both are critical for mitochondrial quality control [45, 50]. To determine whether mitochondrial dysfunction underlies the exacerbated double-mutant phenotype, we measured mitochondrial membrane potential (MMP) in WT, *sod2*-M1, *atg43*Δ, and *sod2*-M1 *atg43*Δ strains using MitoTracker staining. In WT cells, mitochondrial fluorescence decreased significantly during quiescence (**Fig. 5D**), indicating a reduction in MMP, a phenomenon consistent with reports in hematopoietic stem cells [51, 52]. In the *sod2*-M1, *atg43*Δ, and *sod2*-M1 *atg43*Δ mutants, MitoTracker signals during quiescence were all elevated above WT levels. Crucially, the signal in the double mutant was significantly higher than in either single mutant (**Fig. 5E**). This indicates that autophagy deficiency and disrupted *sod2* APA independently cause abnormal MMP, and their combination produces a synergistic effect, suggesting exacerbated mitochondrial dysfunction.

Collectively, these findings demonstrate that APA regulation of *sod2* becomes indispensable under autophagy-impaired conditions. APA-mediated regulation of *sod2* likely safeguards long-term cell survival during quiescence by maintaining mitochondrial homeostasis.

### Pabp is an upstream regulator of *sod2* APA

We next sought to identify the factors regulating *sod2* APA. To this end, we focused on 145 known trans-acting APA regulators (**Tab. S2**) and calculated their correlation with both the PDUI and expression levels of *sod2* across our 15 time points. Trans-factors regulating *sod2* APA can be classified into two categories: (1) positive regulators of 3’ UTR length, whose expression positively correlates with PDUI (**Fig. 6A top**); and (2) negative regulators of 3’ UTR length, whose expression negatively correlates with PDUI (**Fig. 6A bottom**). Through this analysis, we identified one potential positive regulator and 26 potential negative regulators of *sod2* APA (**Fig. 6B**).

**Fig. 6.**
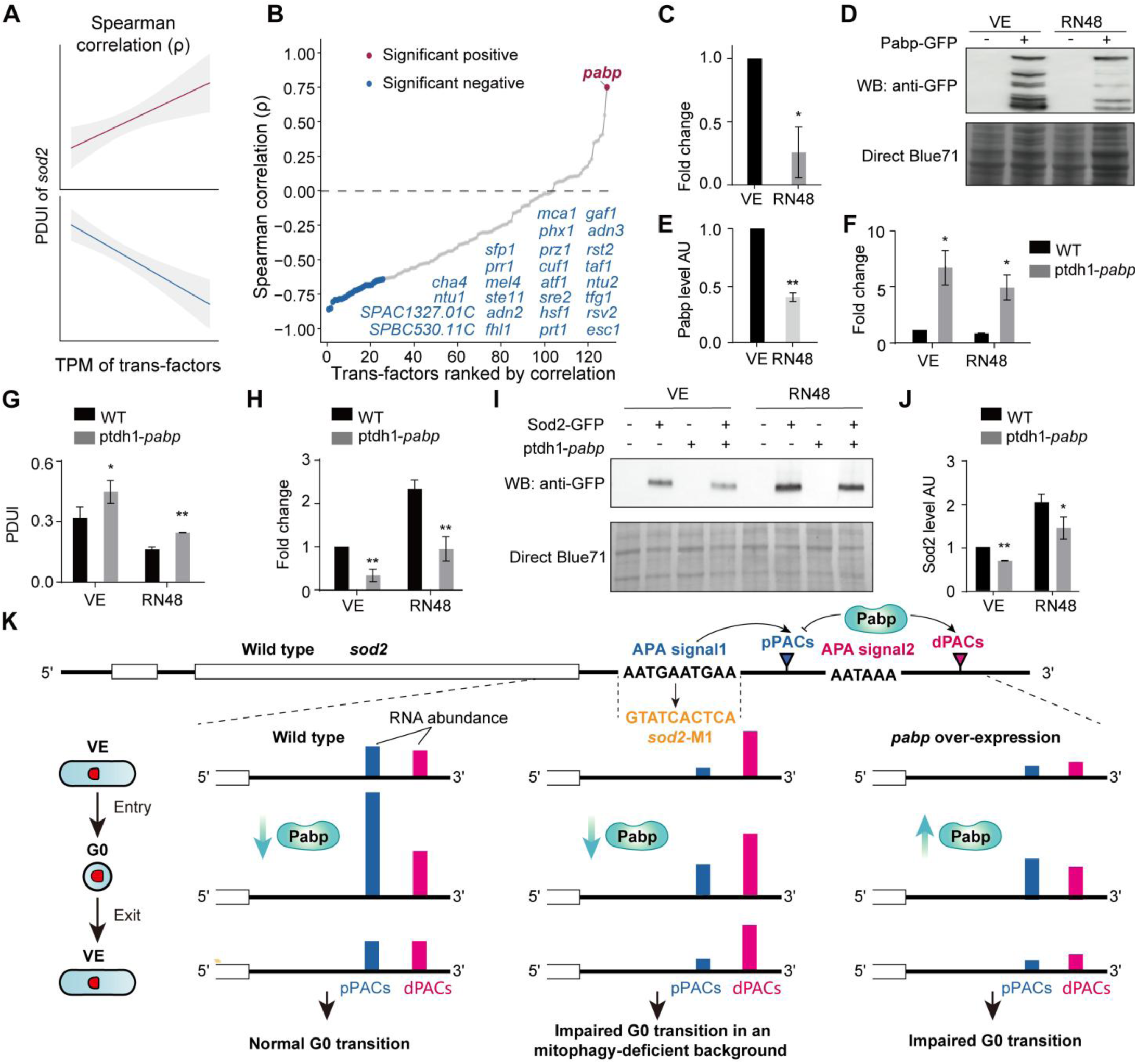
*pabp* is an upstream regulator of *sod2* APA. **(A)** Schematic illustrating the definition of positive and negative regulation of *sod2* by trans-factors. **(B)** Distribution of spearman correlation values of all trans-factors, with significant genes marked. (**C**) Endogenous *pabp* expression in wild-type (WT) cells during vegetative growth (VE) and quiescence after 48 h of nitrogen starvation (RN48). Expression level normalized to VE. (**D**) Western blotting of Pabp-GFP in WT VE cells and RN48 cells. The parental strain (no tag) was employed as a negative control. The protein loading was confirmed by Direct Blue 71 staining. (**E**) The Pabp-GFP band intensity was characterized via three independent western blotting results, and the mean ± S.D. was presented. AU: arbitrary unit. (**F**) *pabp* overexpression driven by the *tdh1* promoter during vegetative growth and quiescence. Expression level normalized to WT. (**G**) PDUI of *sod2* in WT and ptdh1-*pabp* strains. **(H)** *sod2* expression in WT and ptdh1-*pabp* strains during vegetative growth and quiescence. Expression level normalized to WT. (**I**) Western blotting of Sod2-GFP in wild-type and ptdh1-*pabp* cells under VE and RN48 conditions. Protein loading was confirmed by Direct Blue 71 staining. **(J)** The Sod2-GFP band intensity was characterized via three independent western blotting results, and the mean ± S.D. was presented. (**K**) Model for APA-mediated *sod2* expression regulation during quiescence transition. In WT cells, Pabp downregulation upon G0 entry promotes APA switching from dPAS (long 3’ UTR) to pPAS (short 3’ UTR), enhancing *sod2* mRNA levels; the pattern reverts upon G0 exit. Pabp overexpression inhibited the APA switch from dPAS to pPAS, confirming its role as a negative regulator of pPAS usage. In the *sod2*-M1 mutant, pPAS disruption prevents this switch, leading to impaired Sod2 expression and compromised quiescence fitness under mitophagy-deficient stress. Pabp thus acts as a key upstream regulator of *sod2* APA during quiescence transition. *\* P* < 0.05, *\*\*P* < 0.01. AU: arbitrary unit.

Among these, *pabp* was the only significantly positive regulator and was therefore selected for further analysis and validation. Pabp is the *S. pombe* ortholog of mammalian poly(A)-binding protein PABPC1 [53]. In human cells, PABPC1 has been shown to regulate APA site selection: high expression promotes the usage of distal APA sites (increasing PDUI and lengthening 3’ UTRs), whereas low expression favors proximal APA sites (decreasing PDUI and shortening 3’ UTRs) [54–56]. Thus, Pabp acts as a negative regulator of proximal APA sites (or a positive regulator of distal APA sites). However, its function during quiescence transitions remains elusive, and there is currently no evidence regarding whether Pabp regulates APA site selection at *sod2* during this process.

If the function of Pabp is conserved between mammals and fission yeast, genes regulated by Pabp should preferentially utilize distal sites (increasing PDUI and 3’ UTR length) when Pabp is upregulated, and proximal sites (decreasing PDUI and 3’ UTR length) when Pabp is downregulated. Our experimental data supported this hypothesis: at RN48 of quiescence entry, *pabp* RNA expression was significantly downregulated compared to the VE stage (**Fig. 6C**), and this downregulation was also observed at the protein level by western blotting (**Figs. 6D and 6E**). Consistent with reduced Pabp abundance, *sod2* indeed showed a preference for the proximal site (decreased PDUI; **Fig. 4B**). This indicates that the relationship between *sod2* APA selection and Pabp expression in *S. pombe* is consistent with the mechanism of Pabp-mediated APA regulation observed in mammals.

However, the above observation only establishes a correlation between *sod2* APA dynamics and *pabp* expression, not causality. To address this, we constructed a *pabp* overexpression strain (see **MATERIALS AND METHODS**) to determine whether elevated Pabp levels would induce the expected shift in *sod2* APA site selection. Indeed, upon *pabp* overexpression in both VE and quiescent cells (**Fig. 6F**), the PDUI of *sod2* increased significantly compared to the WT (**Fig. 6G**). Furthermore, due to the regulatory effect of *sod2* APA on its own mRNA abundance, we also observed a significant reduction in *sod2* mRNA abundance (**Fig. 6H**). Accordingly, Sod2 protein levels were also diminished in the pabp overexpression strain under both vegetative growth and quiescent conditions (**Fig. 6I and 6J**).

Flow cytometry analysis revealed that after 48 hours of nitrogen starvation, only 39.4% of *pabp*-overexpressing cells were at the 1C DNA content stage. Moreover, after two weeks of nitrogen starvation followed 12 hours after nitrogen replenishment, these cells remained arrested at the 1C stage and failed to recover to the 2C state (**Fig. S7A**). Trypan blue exclusion assays showed that by the third week of nitrogen starvation, the viability of the *pabp* overexpression strain was significantly lower than that of the WT (**Fig. S7B**). Colony formation assays further demonstrated that the colony-forming ability of the *pabp* overexpression strain was markedly impaired during both the proliferative and quiescent phases; after two weeks of nitrogen starvation, this strain nearly completely lost its ability to form colonies (**Fig. S7C**). These results indicate that Pabp regulates quiescence transitions. It is worth noting, however, that while *pabp* overexpression affects quiescence transitions, this effect may not be mediated solely through *sod2* APA, as Pabp regulates the APA of many other genes [57, 58].

Collectively, these findings demonstrate that Pabp, as a positive regulator of distal APA site selection, is downregulated during quiescence entry, thereby promoting decreased distal site usage, reduced PDUI, and 3’ UTR shortening of *sod2*. This shortening of the *sod2* 3’ UTR contributes to the upregulation of its own expression.

## DISCUSSION

Quiescence is a highly conserved and vital survival strategy evolved by organisms. Although numerous studies have investigated quiescence [22, 59–61], the regulatory mechanisms governing entry into and exit from this state remain poorly elucidated, particularly regarding the role of post-transcriptional regulation. APA, a core post-transcriptional regulatory mechanism [18, 62], has rarely been explored in the context of quiescence transitions. In this study, we systematically investigated the dynamic landscape of APA during quiescence transitions in the model organism *S. pombe*, and explored the molecular mechanisms by which APA regulates gene expression through 3’ UTR length alterations to influence quiescence. Using 3’-end mRNA sequencing, we constructed a genome-wide atlas of APA dynamics during *S. pombe* quiescence transitions (**Figs. 1 and 2**) and revealed that APA-mediated 3’ UTR remodeling potentially regulates gene expression during these transitions (**Fig. 3**). Specifically, using *sod2* as a representative example, we systematically elucidated the role and mechanism of APA in regulating quiescence (**Fig. 6G**): during quiescence entry, *sod2* preferentially utilizes proximal poly(A) sites, resulting in shorter 3’ UTRs; conversely, during quiescence exit or vegetative growth, it favors distal poly(A) sites and longer 3’ UTRs. This APA-mediated regulation of *sod2* 3’ UTR length modulates *sod2* expression levels during quiescence entry, thereby affecting quiescence transitions under autophagy-deficient conditions. Furthermore, Pabp was identified as an upstream positive regulator of distal APA site usage of *sod2*. Our study delineates a Pabp-APA-*sod2* regulatory axis during quiescence transitions, highlighting the critical role of post-transcriptional regulation in quiescence adaptation and stress tolerance, and providing an important case study for understanding the post-transcriptional control of cellular quiescence transitions.

During quiescence transitions, we observed genome-wide APA remodeling (**Figs. 2, 3C, and 3D**). This global remodeling exhibited distinct differences between phases, such as between quiescence entry and VE, or between quiescence exit and the fully quiescent state (RN48). Within individual phases, changes across time points appeared relatively subtle, such as during the first 8 time points of entry or the last 4 time points of exit. However, dramatic shifts occurred at critical transition points, such as at 0.5 hours of nitrogen starvation (RN0.5) versus VE, or at 1 hour of nitrogen replenishment (AN1) versus the fully quiescent state (RN48). This pattern suggests that global APA remodeling during quiescence transitions is not a continuous, gradual process, but rather manifests as a rapid, “switch-like” or pulsatile regulation. This characteristic likely reflects a stress adaptation strategy in response to drastic environmental changes, where intense alterations are triggered over a short period, followed by the establishment of a new homeostatic balance [63, 64]. Interestingly, at 48 hours of nitrogen starvation (RN48), a massive APA shift occurred compared to the previous time point (RN24). Further investigation with denser sampling between RN24 and RN48 hours will be required to fully explain this dramatic transition.

Although APA-induced 3’ UTR changes involve a considerable number of genes during quiescence transitions (**Fig. 3C**), the biological significance of these changes remains debatable. First, the magnitude of 3’ UTR length alteration for these genes is relatively small, averaging between 8 and 18 nt (**Fig. 3D**). Such narrow ranges may not necessarily contain functional regulatory elements, suggesting that most APA events might be unrelated to gene expression regulation. Second, integrated analysis of 3’ UTR changes and gene expression variations identified only 29 significantly correlated genes (**Fig. 3I**), further indicating that APA-mediated gene regulation is not a widespread phenomenon. This observation is consistent with the previous findings across multiple species and cell lines, where APA differences and gene expression differences were largely decoupled [65], and aligns with the molecular error hypothesis of APA [32]. Clearly, APA may possess other potential functions independent of mRNA abundance, although these fall outside the scope of the current study.

Despite the fact that functional APA events may represent only a minority, existing cases demonstrate that APA can serve as an important strategy for gene expression regulation in specific contexts [18]. In this study, we newly demonstrated that 3’ UTR shortening of *sod2* during quiescence enhances *sod2* expression, representing another typical example of APA regulation. Notably, although APA regulates *sod2* expression, this regulation appears to be context-dependent: we observed this regulatory effect during nitrogen-deprivation-induced quiescence entry, but not during the VE stage or quiescence exit (**Fig. 4E**). The specificity of this regulation likely depends on extrinsic trans-acting factors, which warrants further investigation.

Furthermore, our data revealed that *sod2* expression levels during quiescence entry were significantly higher than those at the VE stage or during quiescence exit. Notably, this magnitude of upregulation was substantially greater than the difference observed between *sod2*-M1 and WT (**Fig. 4E**). This observation suggests that, in addition to post-transcriptional regulation via APA, *sod2* expression during quiescence transitions is also likely governed by other regulatory mechanisms, such as transcriptional initiation. Indeed, previous studies in other species have demonstrated that *sod2* is regulated at the transcriptional level. For instance, in mammalian cells, AKT is inactivated during quiescence, leading to the activation of FOXO3a. Active FOXO3a binds to the *sod2* promoter to drive its transcription [66, 67]. Conversely, upon entry into the proliferative phase, AKT activation results in FOXO3a inactivation, thereby abolishing its ability to drive or enhance *sod2* transcription [66]. Collectively, these findings and our results together indicate that *sod2* is subjected to both transcriptional and post-transcriptional regulation during quiescence transitions. While the present study focused on elucidating the post-transcriptional APA-mediated regulation of *sod2*, the specific transcriptional regulatory mechanisms in *S. pombe* remain to be clarified in future investigations.

Notably, the *sod2* APA single mutant (*sod2*-M1) did not exhibit obvious phenotypic defects during nitrogen starvation-induced quiescence maintenance. This can be understood from three perspectives. First, the regulatory amplitude of APA on *sod2* expression may be limited (**Fig. 4E**); the reduction in *sod2* expression likely only depletes the redundant pool of Sod2, thereby failing to elicit obvious quiescence-related phenotypes. Second, when *Sod2* expression levels are moderately reduced, compensatory mechanisms by other antioxidant enzymes (e.g., Sod1, catalase) may suffice to maintain redox homeostasis. Third, nitrogen starvation in the laboratory represents a single stressor, whereas *S. pombe* in natural environments faces multiple concurrent pressures, including temperature fluctuations, osmotic stress, oxidative stress, nutrient deprivation, and interspecies competition. We speculate that the moderate upregulation of *Sod2* resulting from *sod2* 3’ UTR shortening may be crucial for cell survival under multi-stress conditions. Supporting this notion, *sod2* APA proved functionally essential under autophagy-deficient backgrounds. Therefore, the absence of a phenotype under laboratory conditions does not equate to a lack of function, underscoring the need to re-evaluate APA contributions under a broader range of conditions.

Beyond *sod2*, numerous other genes exhibited coupled changes in APA and gene expression (**Fig. 3**). Many of these genes have been implicated in quiescence regulation through high-throughput screens [37, 68], such as *vac7* and *hul6*, which have been experimentally confirmed to regulate cellular quiescence. Our study also identified many genes not previously associated with quiescence, and the regulatory role of APA on their expression remains largely unexplored, warranting further investigation (**Fig. 3K**).

## MATERIALS AND METHODS

### Media and Strains

Complete medium (YES), complete medium supplemented with 5-fluoroorotic acid (+FOA), minimal medium (EMM), minimal medium without nitrogen (EMM-N), MSL lacking uracil (MSL-Ura), and sporulating medium (SPA) were utilized [69]. The *S. pombe* strains used in this study are listed in **Tab. S3**. GFP-tagging, deletion, and *pabp* overexpression strains were generated by PCR-based homologous recombination as described [70]. The DNA fragments for gene deletion or tagging contained drug-risistant genes kanMX6, natMX6. The strains were screened by YES plate with G418 or zhongshengmycin. Specific point mutation *sod2*-M1 was introduced using the “pop-in, pop-out” allele replacement method [71]. Strains overexpressing *pabp* from the endogenous *pabp* locus under the control of the *tdh1* promoter were generated. Random spore analysis was applied by following the method described by Bahler et al.[70]. Primers and plasmids used in this study were listed in **Tabs. S4, and S5**, respectively.

### G0 induction and survival assay

Heterothallic (*h*^-S)^ prototroph fission yeast cells were cultivated at 30°C for 24h. The cells were then cultured in EMM for 12 h and subcultured twice to remove dead cells. Subsequently, cells were collected, washed twice with sterile water, and resuspended in EMM-N to induce G0 entry. The initial OD₆₀₀ was adjusted to 0.3–0.5. Quiescent cells were cultured at 30°C with shaking at 220 rpm.

At indicated time points (0, 1, 7, 14, 21, and 28 days), a 90 μL aliquot of the culture was mixed with 10 μL of trypan blue and incubated for 3 min at room temperature. The percentage of trypan blue-negative cells was defined as the proportion of viable cells. At the same time points, serially diluted (10-fold serial dilutions), and spotted onto YES plates. The plates were incubated at 30°C for 3 days, and the number of colonies formed was counted. Colony formation efficiency for detecting the MC was calculated as the percentage of colonies formed relative to the number of cells plated.

### RT-qPCR

Total RNA was isolated using the MasterPure™ Yeast RNA Purification Kit (MPY03100, Epicentre). RNA samples contaminated with DNA were treated with TURBO DNase (AM2238, Invitrogen). Complementary DNA (cDNA) was synthesized using the HiScript III RT SuperMix for qPCR (R323-01, Vazyme). Quantitative PCR (qPCR) was performed with the ChamQ Universal SYBR qPCR Master Mix (Q711-02, Vazyme) on a LightCycler 480 instrument (Roche). The primers used for RT-qPCR are listed in **Tab. S4**.

### Microscopic analysis

Fluorescence microscopy were performed using a Zeiss Axio Imager Z2 microscope (Carl Zeiss MicroImaging). Raw images were processed with ZEN lite 2012 (Carl Zeiss MicroImaging). mitochondrial potential was measured as described previously [72], with minor modifications. Briefly, fission yeast cells in logarithmic growth phase and G0 cells after 48 h of nitrogen starvation were washed once with their respective media: EMM for log-phase cells and EMM-N for G0 cells. To assess functional mitochondria, washed cells were resuspended in medium containing 100 nM MitoTracker (Invitrogen Corporation, Carlsbad, CA, USA) and incubated for 30 min in the dark at room temperature. After staining, cells were washed three times to remove excess probes. Fluorescence was measured at excitation/emission wavelengths of 579/599 nm for MitoTracker.

### Western blotting

Protein samples for western blotting were isolated as previously described [25]. Cells were harvested at the indicated time points and were lysed by bead-beating and extracted in lysis buffer [100 mM ammonium bicarbonate, 8 M urea, and - protease inhibitors (P88300, ABCone)]. The protein samples were mixd with equal volume of 2×SDS loading buffer [(60 mM Tris-HCl, pH6.8), 20% glycerol, 4% SDS, 0.04% bromophenol blue, 5% β-mercaptoehanol] and were heated at 98°C for 5 minutes. 10% SDS-PAGE gels were used for separating proteins. Anti-GFP (clones 7.1 and 13.1, Roche) antibodies were utilized for probing the Sod2-GFP. Western blotting was conducted independently on at three times to ensure reproducibility. Protein loading was assessed using Direct Blue 71 staining [73].

### FACS

Collect cells at the indicated time points (OD_600_ = 0.3) by centrifuging. Resuspend with 300 μl of of sterilized water, add 700 μl of 95% ethanol while vortex. Fix at 4 °C overnight. 3,000 rpm 2min, remove supernatant. Wash with 1 ml of 50mM sodium citrate 3 times. Add 1 ml of 50mM sodium citrate to suspend the yeast, add 25μl 10 mg/ml RNaseA, 37°C 6h. Add 1ml of 25 μg/ml PI solution, stain at 4 °C overnight. Sonicate 30s, filter the sample and load on the flow cytometry. The area of PE channel is regarded as DNA level.

### 3’-end mRNA-seq

3’ mRNA-seq libraries were prepared with 500 ng of total RNA using the Quant-seq 3’ mRNA-seq V2 Library Prep Kit REV (Lexogen) according to the manufacturer’s instructions. Briefly, 500 ng total RNA was used as starting material. Poly(A) containing RNA was reverse transcribed by anchored oligo(dT) primer, followed by RNA removal and second-strand cDNA synthesis with random primers containing part of the Illumina adaptor sequence. PCR amplification was then performed to get the libraries. The completed libraries underwent quality control using a Bioanalyzer (Agilent) with the High Sensitivity DNA assay. Library concentrations were measured using a Qubit 1× dsDNA HS Assay Kit (Thermo Fisher Scientific). Deep sequencing was performed using the NovaSeq6000 platform (Illumina) and 150-bp reads.

### General processing of APA data and PAC identification

To obtain more accurate annotations of 5’ UTR and 3’ UTR regions, annotated gene termini were further refined using public RNA-seq datasets (SRR27030748, SRR26525761, SRR25915681, SRR23685324, SRR32068752 and SRR24996077). UTR extension was determined based on continuous RNA-seq read coverage and was constrained to avoid overlap with neighboring genes while maintaining biologically reasonable UTR annotations across the genome.

Only R1 reads in paired-end sequencing were used for downstream analyses. Adaptor trimming was performed with cutadapt [74], followed by quality filtering with fastp [75] to remove low-quality and short reads. The processed reads were then mapped to the *S. pombe* genome ASM294v2 using STAR v2016-03-14 [76], and alignment files were processed with SAMtools v1.3 [77]. Only uniquely mapped reads were retained for subsequent analyses and the 3’ most site of each mapped read was annotated as the PAS. To eliminate potential internal priming artifacts, reads mapped to adenine-rich genomic regions containing at least six consecutive adenines within a 10 nt sliding window were excluded from further analysis [78].

To minimize false-positive PAS identification caused by sequencing noise, we applied a Poisson distribution-based model to assess read enrichment at each candidate PAS. Candidate PASs with read counts significantly exceeding the expected background level were considered high confidence sites. Based on this model and the ultra-high sequencing depth of our dataset, a minimum read-count cutoff of 4 was determined. Accordingly, only PASs supported by at least 4 reads were retained for downstream analyses across all environments.

To determine the optimal clustering distance for grouping nearby PAS, we follewed the threshold of 7 used by previous study [20]. The QuantifyPoly(A) method [79] was then used to cluster PASs: PASs separated by ≤7 nt were grouped into the same PAC, whereas those separated by >7 nt were assigned to distinct PACs. Within each PAC, the PAS supported by the largest number of reads was designated as the representative site.

### 3’ RACE experimental validation of APA events

To validate the authenticity of detected APA events in this study, 3’ RACE was performed using 3’-Full RACE Core Set with PrimeScript™ RTase (6106, TakaraBio Technology, China) according to the manufacturer’s instructions. Briefly, 1 μg of total RNA was reversely transcribed into cDNA using 3’ RACE adaptor. Then the cDNA of a specific gene was amplified using 3’ RACE outer primer and gene specific outer primer, followed by a nested PCR using 3’ RACE inner primer and gene specific inner primer. The sequences of all gene-specific primers were described in **Tab. S4**.

### Differently expression gene analysis

Differential gene expression analysis was performed using the DESeq2 and APAdeg [65, 80] package in R. Raw read counts for each gene were used as input, and normalization was carried out using the default median-of-ratios method implemented in DESeq2. Genes with an adjusted *P* value (Benjamini-Hochberg correction) < 0.05 and an absolute log2 fold change ≥ 1 were considered significantly differentially expressed, where fold change is defined as the expression in RN points divided by the expression in VE, or the expression in AN point divided by the expression in RN48.

### Down-sampling

To minimize the potential impact of differences in sequencing depth in the comparison across samples, PAC read counts were normalized by down-sampling each sample to the lowest sequencing depth observed across all samples (15 time points). For samples with higher sequencing depth, reads were randomly subsampled without replacement until the total number of reads matched that of the sample with the lowest sequencing depth. Samples with the lowest sequencing depth were retained unchanged.

### Quantification of 3’ UTR length in genes

In many cases, multiple PACs were concurrently used within a gene, generating transcripts with different 3’ UTR lengths. Therefore, using a single PAC to represent the 3’ UTR length cannot fully capture the heterogeneity of polyadenylation. We thus calculated the abundance-weighted 3’ UTR length of a gene from all its isoforms based on the formula:

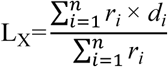

where *n* is the total number of PACs identified for gene X, *r_i_* denotes the number of reads supporting the *i*-th PAC, and *di* is the 3’ UTR length of transcript generated from the *i*-th PAC.

### Quantification of distal APA usage in genes

Distal poly(A) site usage for a gene was quantified using the Percentage of Distal poly(A) site Usage Index (PDUI). Briefly, by reconstructing APA site data into RNA coverage profiles, we defined the boundary (i.e., breakpoint) separating proximal and distal APA sites for a gene across time points **(Fig. S3A)**. PDUI was calculated as the fraction of read counts assigned to distal PACs among the total read counts of all PACs within the gene in each time point, as the following formula:

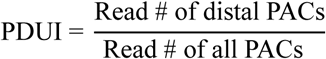

Statistical significance of PDUI changes was assessed for each gene using a two-tailed Fisher’s exact test based on proximal and distal PAC read counts. *P* values were adjusted for multiple testing using the Benjamini-Hochberg method. Genes with an adjusted *P* value < 0.05 were considered to exhibit significant poly(A) site switching.

### Nucleotide profile and motif analysis

Nucleotide profiles in the flanking regions of PACs were analyzed at the PAC abundance-weighted level, where each PAC was weighted by its read abundance. De novo 6-mer motif discovery was performed using the findMotifsGenome.pl program implemented in the HOMER software package [81]. Motif enrichment was analyzed in three predefined regions surrounding the poly(A) site (−60 to −31 nt, −30 to −1 nt, and 0 to +30 nt), as well as within the full 3’ UTR region of genes. *P* values based on the Fisher’s exact test were used to indicate the significance of enrichment.

### Gene ontology enrichment analysis

GO enrichment analysis was performed using the clusterProfiler [82] package in R. All protein coding genes detected across the 15 time points were used as the background gene set. Statistical significance and odds ratios were calculated using Fisher’s exact test. Because some gene sets contained relatively few genes, multiple-testing correction could potentially exclude biologically relevant terms. Therefore, GO terms with *P* < 0.05 were considered putatively enriched.

### Statistical tests

All statistical analyses in this study were performed using the R programming language. The details of the statistical methods for each figure are described in the corresponding figure legends.

## Data availability

Sequence data that support the findings of this study have been deposited in the National Genomics Data Center under the accession number PRJCA071096 (https://ngdc.cncb.ac.cn/gsa/s/fdE5k2Tk). The authors declare that all other data supporting the findings of this study are available within the article and its supplementary files.

## ACKNOWLEDGMENTS

We are grateful to Xiangzhou Meng at Shanghai Jiao Tong University and Tomoyasu Sugiyama at ShanghaiTech University for kind help in wet experiments.

## FUNDINGS

This study was supported by National Natural Science Foundation of China (32270704, 32472630, and 32670917), National Science and Technology Innovation 2030 Major Projects for ‘Brain Science and Brain-Inspired Research’ (2022ZD0214400), and Medical-Engineering Crossover Fund of Shanghai Jiao Tong University (YG2025QNB51).

## AUTHOR CONTRIBUTIONS

C. X. conceived and designed the study. C. X., X. D., and L. Z. wrote the paper. L. Z., X. D., C. X., and Y. J. analyzed the data. X. D., Z. D., and L. G. did the wet experiments. B. S. provided the pipeline of APA analysis. T. Z., L. L., S. H., and Y. Z. joined the discussions.

## CONFLICT OF INTEREST STATEMENT

The authors declare no conflict of interest.

## Supplementary figures and legends

**Fig. S1.**
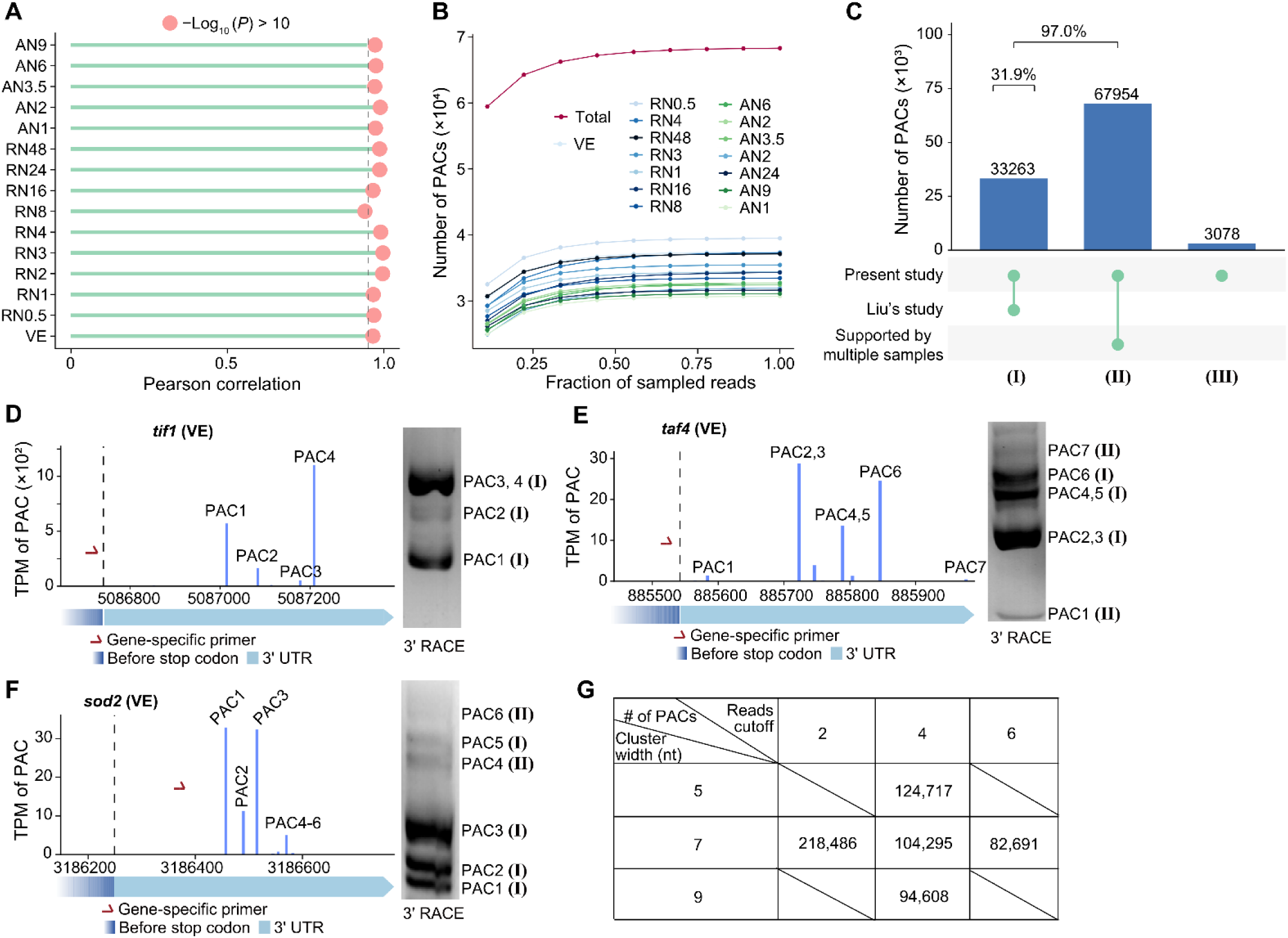
Quality control and characterization of PACs. (**A**) Pearson correlations between biological replicates for PAC expression levels in each time point. (**B**) Saturation analysis of PAC detection in individual and combined time points. (**C**) Venn diagram showing the overlap of PACs identified in the present study and Liu’s study, and supported by multiple samples. *tif1* (**D**), *taf4* **(E)** and *sod2* **(F)** as examples for 3’ RACE validation of PACs identified by Quant-seq analysis. (**G**) Number of PACs identified under different cut-offs of cluster distances and minimum read counts.

**Fig. S2.**
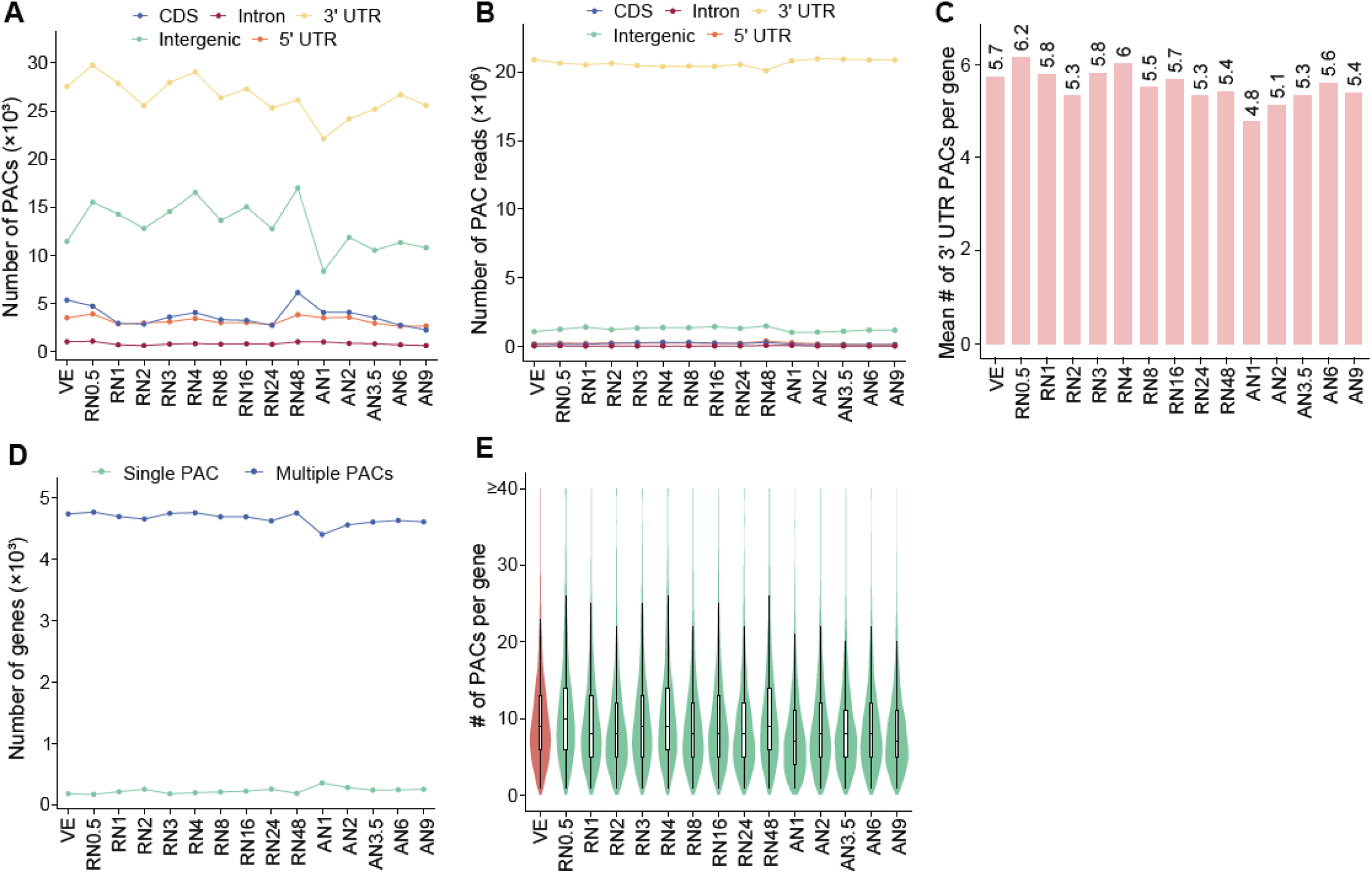
APA characteristics among 15 time points. (**A**) Distribution of PAC counts across different genomic regions in various time points. (**B**) Distribution of PAC read counts across different genomic regions in various time points. (**C**) Number of 3’ UTR PACs per gene across time points. (**D**) Number of genes with single or multiple PACs across various time points. (**E**) Distribution of gene counts categorized by the number of expressed PACs across time points.

**Fig. S3.**
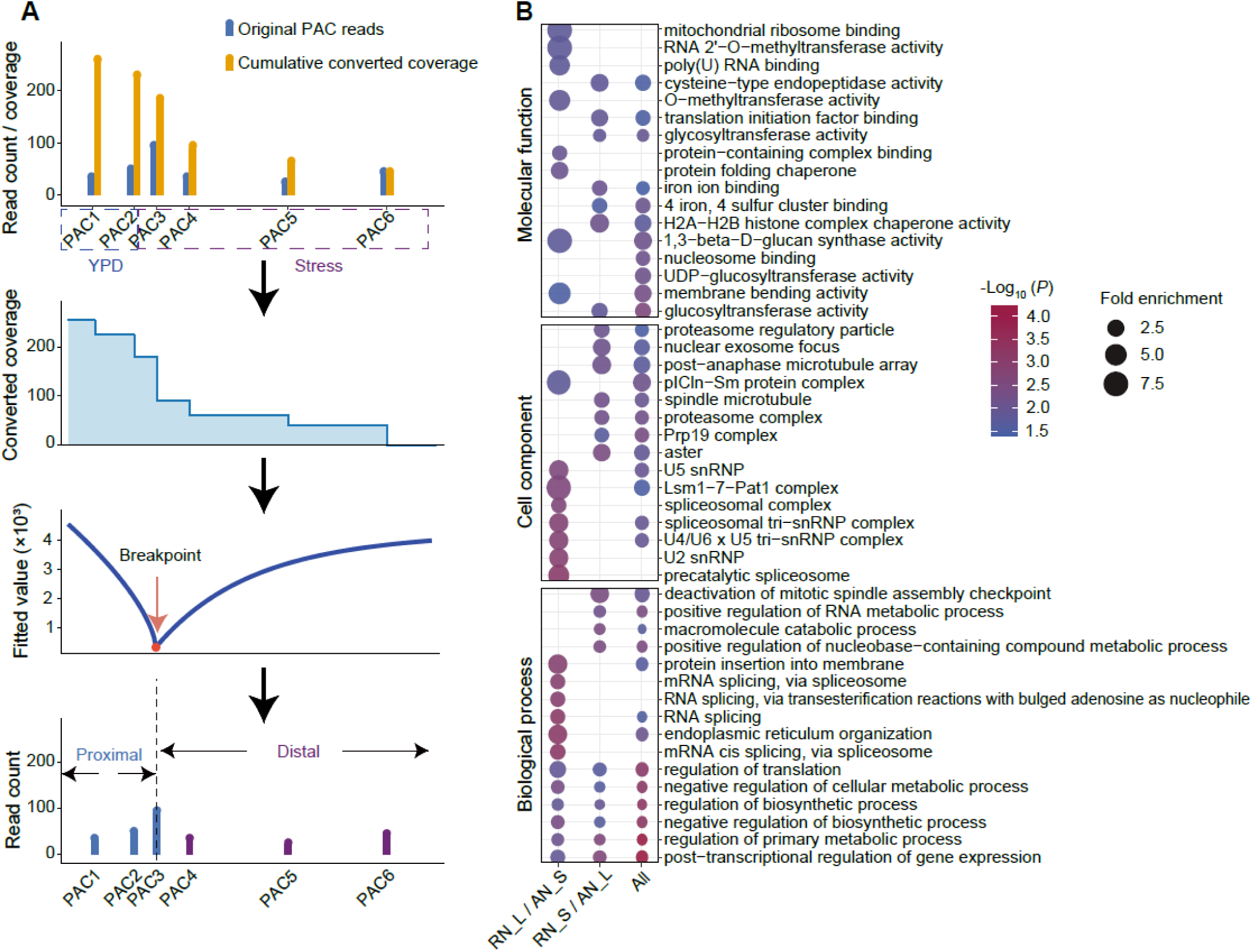
Quantification method and functional characterization of 3’ UTR variability. **(A)** Schematic illustrating the determination of the breakpoint between proximal and distal PACs. Original PAC read counts were converted into cumulative coverage and fitted to identify the breakpoint. PACs upstream and downstream of the breakpoint were classified as proximal and distal PACs, respectively. **(B)** Gene Ontology enrichment analysis of genes with opposite 3’ UTR length changes during quiescence entry and exit. Left: genes with 3’ UTR lengthening upon entry and shortening upon exit (RN_L/AN_S). Middle: genes with 3’ UTR shortening upon entry and lengthening upon exit (RN_S/AN_L). Right: the combined gene set from both groups above (All).

**Fig. S4.**
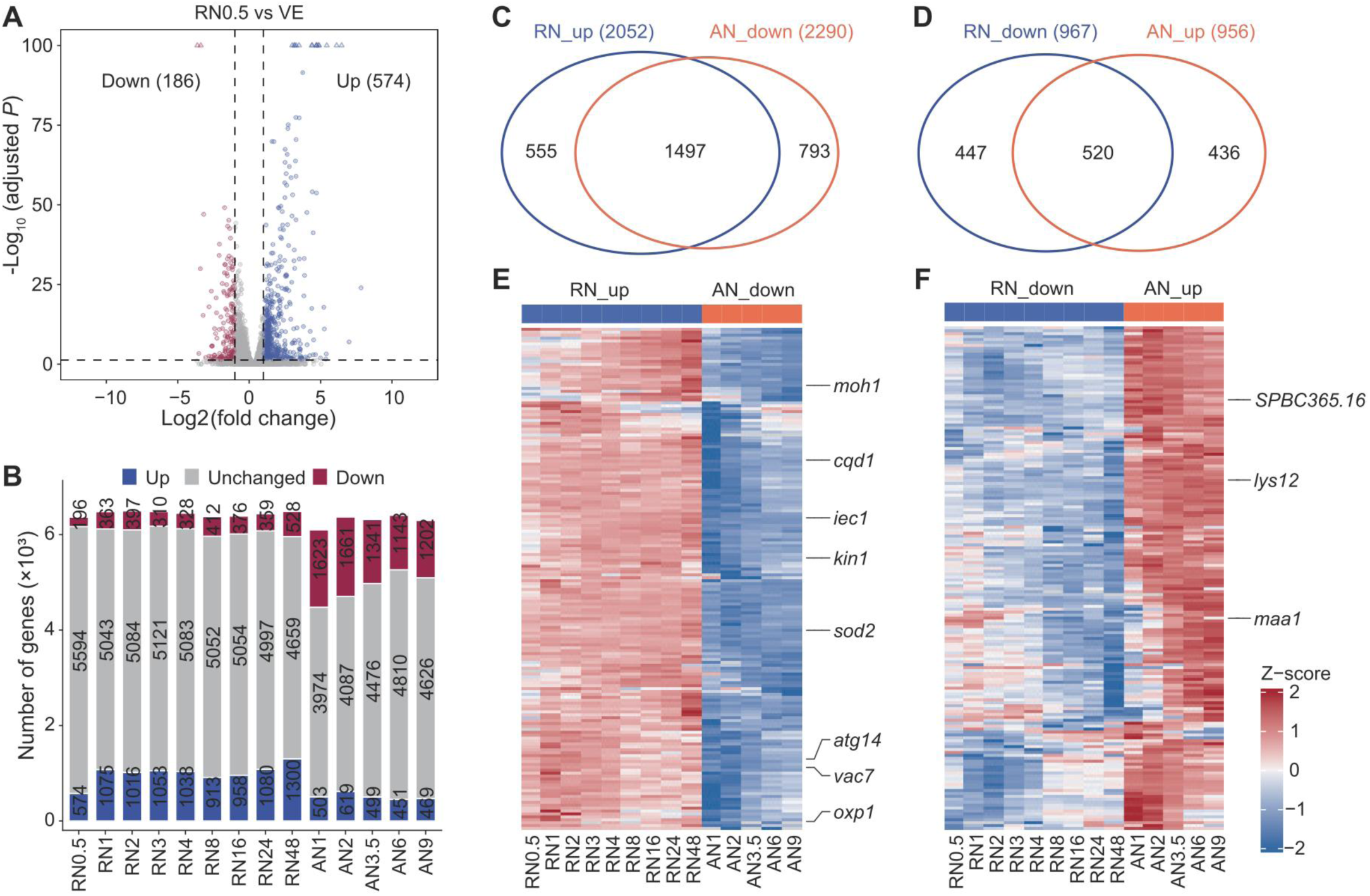
Differently expressed genes (DEG) during quiescence transitions. (**A**) DEGs in RN0.5 as an example. (**B**) Number of DEGs in each time point. (**C**) Overlap of genes up-regulated in quiescence entry stages (RN_up) and down-regulated in quiescence exit stages (AN_down). (**D**) Overlap of genes down-regulated in quiescence entry stages (RN_down) and up-regulated in quiescence exit stages (AN_up). (**E**) Heatmap showing fold change of up-regulated genes during quiescence entry stages (RN_up) and down-regulated during quiescence exit stages (AN_down). (**F**) Heatmap showing fold change of down-regulated genes during quiescence entry (RN_down) and up-regulated during quiescence exit (AN_up). In (**E**) and (**F**), fold changes were calculated relative to VE for quiescence entry and RN48 for quiescence exit. Selected quiescence-related genes are labeled on the right.

**Fig. S5.**
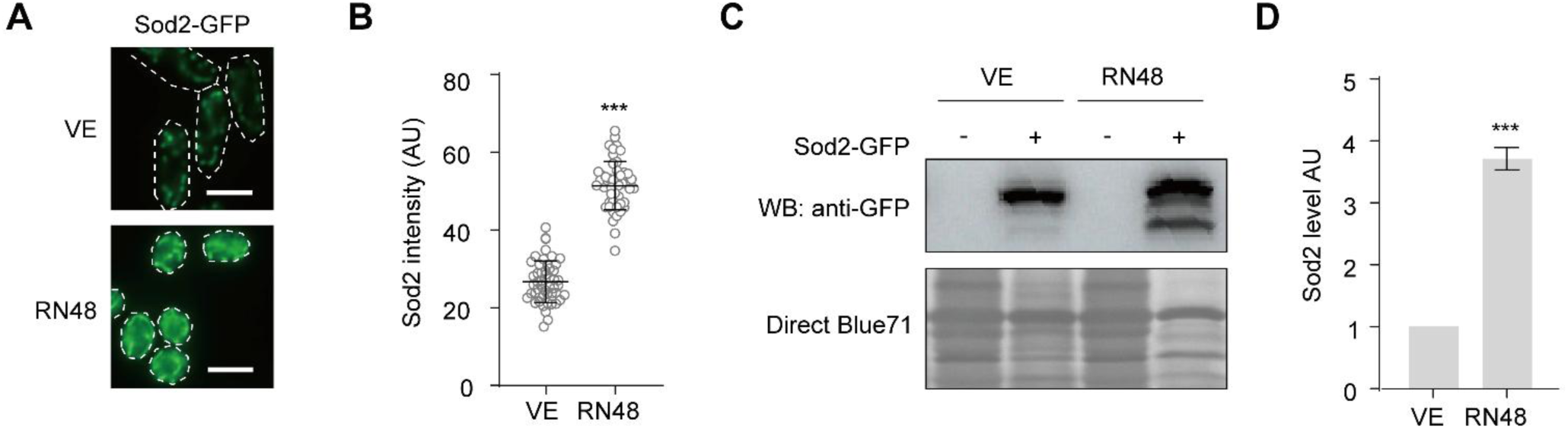
Sod2-GFP is upregulated upon quiescence entry. (**A**) Localization of Sod2-GFP in wild-type (WT) vegetative (VE) cells and quiescent cells after 48 h of nitrogen starvation (RN48). (**B**) A vertical scatter plot of Sod2-GFP signal intensity in wild-type VE cells and RN48 cells. 50 cells were assessed. (**C**) Western blotting of Sod2-GFP in wild type VE cells and RN48h cells. The parental strain (no tag) was employed as a negative control. The protein loading was confirmed by Direct Blue 71 staining. (**D**) The Sod2-GFP band intensity was characterized via three independent western blotting results, and the mean ± S.D. was presented. AU: arbitrary unit. AU: arbitrary unit. *** *P* < 0.001.

**Fig. S6.**
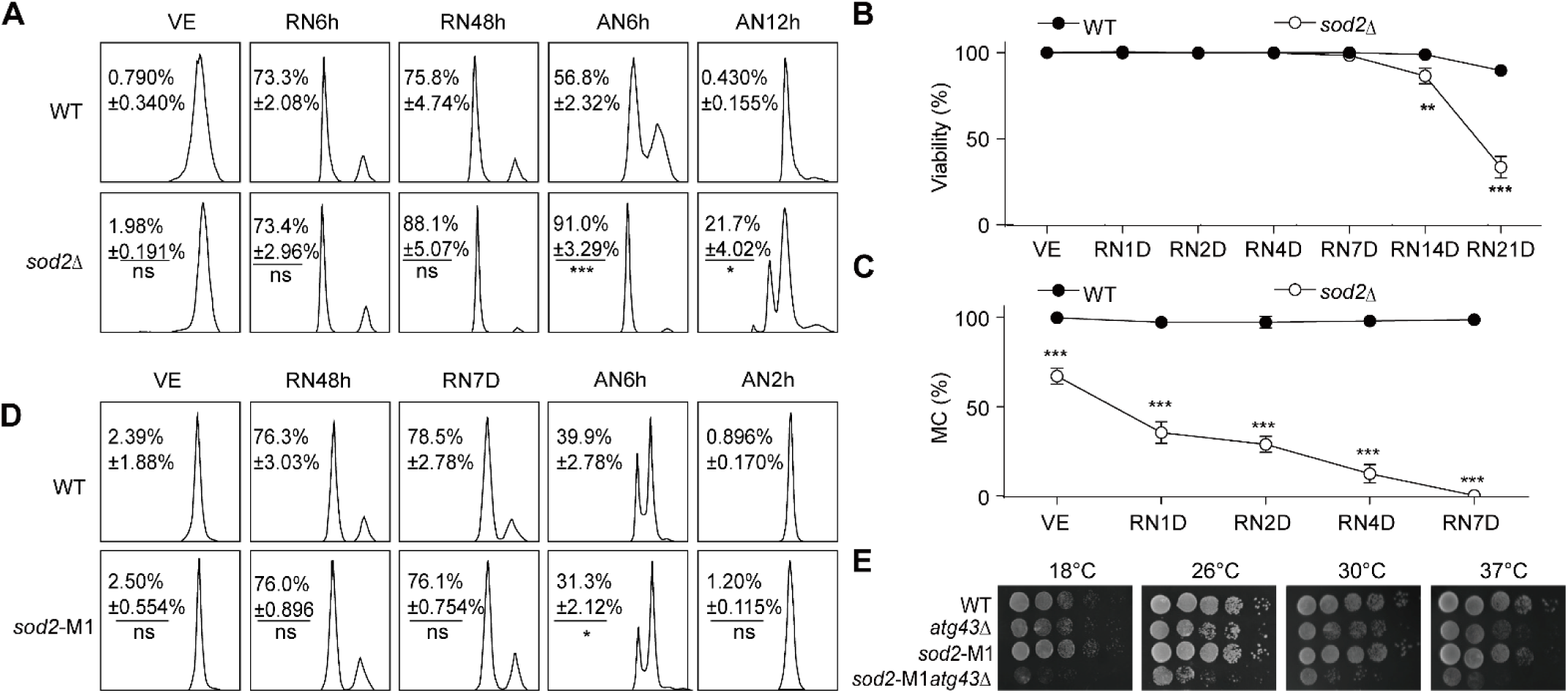
*sod2* is essential for quiescence maintenance and exit. (**A**) Quiescence entry and exit in WT and *sod2*Δ cells were monitored by flow cytometry under nitrogen starvation and subsequent re-addition. Samples were taken during vegetative growth (VE), at 6 h (RN6h) and (RN48h) after nitrogen depletion (for entry), and at 6h (AN6h) and 12h (AN12h) after re-addition (for exit). (**B**) Cell viability in WT and *sod2*Δ cells following nitrogen starvation. Cell viability was assessed by Trypan blue exclusion analysis, examining over 200 cells per experiment, and data are presented as mean ± S.D. from three independent experiments. **(C)** Colony formation assays evaluated mitotic competence (MC) of WT and *sod2*Δ over various time points following nitrogen deprivation. **(D)** Quiescence entry and exit in WT and *sod2*-M1 cells were monitored by flow cytometry under nitrogen starvation and subsequent re-addition. Samples were taken during VE, at 48 h (RN48h) and 7 days (RN7D) after nitrogen depletion and at 6 h (AN6h) and 12 h (AN12h) after nitrogen re-addition. **(E)** Growth of WT, *sod2-*M1, *atg43*Δ and *sod2-*M1*atg43*Δ cells at varying temperatures. Ten-fold serial dilutions were spotted onto complete medium plates and incubated at various temperatures. \*\**P* < 0.01, \*\*\**P* < 0.001.

**Fig. S7.**
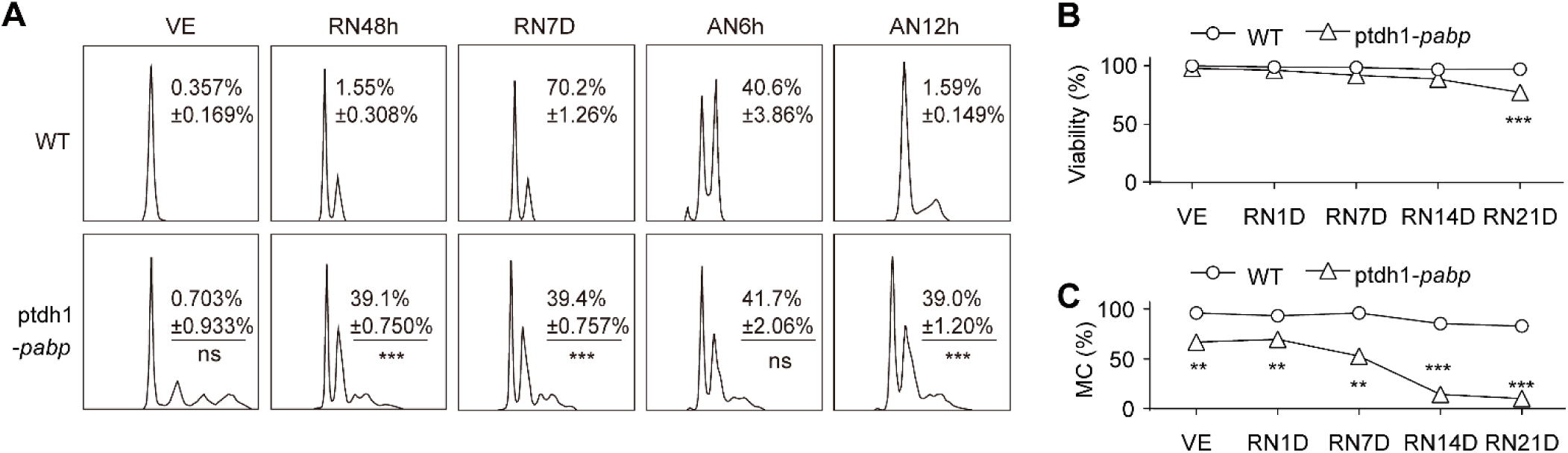
*pabp* overexpression causes pleiotropic defects during quiescence transition. (**A**) Quiescence entry and exit in wild-type (WT) and ptdh1-*pabp* cells upon nitrogen starvation were examined via flow cytometry during vegetative growth (VE), at 48 h (RN48h) and 7 days (RN7D) following nitrogen depletion (G0), and at 6 h (AN6h) and 12 h (AN12h) following nitrogen re-addition. (**B**) Cell viability in WT and ptdh1-*pabp* cells, following nitrogen starvation. Cell viability was assessed by Trypan blue exclusion analysis, examining over 200 cells per experiment, and data are presented as mean ± S.D. from three independent experiments. (**C**) Colony formation assays evaluated mitotic competence (MC) of WT and ptdh1-*pabp* over various time points following nitrogen deprivation. ** *P* < 0.01, *\*\*\* P* < 0.001.

## Supplementary tables and legends

Tab. S1 Summary of Quant-seq 3’ mRNA sequencing data in this study.

Tab. S2 Trans-factors related to APA.

Tab. S3 Strains used in this study, related to Experimental Procedures.

Tab. S4 Primers used in this study.

Tab. S5 Plasmids used in this study.

## Notes

### Competing Interest Statement

The authors have declared no competing interest.

